# Visual motion processing recruits regions selective for auditory motion in early deaf individuals

**DOI:** 10.1101/2020.11.27.401489

**Authors:** Stefania Benetti, Joshua Zonca, Ambra Ferrari, Mohamed Rezk, Giuseppe Rabini, Olivier Collignon

## Abstract

In early deaf individuals, the auditory deprived temporal brain regions become engaged in visual processing. In our study we tested further the hypothesis that intrinsic functional specialization guides the expression of cross-modal responses in the deprived auditory cortex. We used functional MRI to characterize the brain response to horizontal, radial and stochastic visual motion in early deaf and hearing individuals matched for the use of oral or sign language. Visual motion showed enhanced response in the ‘deaf’ mid-lateral *planum temporale*, a region selective to auditory motion as demonstrated by a separate auditory motion localizer in hearing people. Moreover, multivariate pattern analysis revealed that this reorganized temporal region showed enhanced decoding of motion categories in the deaf group, while visual motion-selective region hMT+/V5 showed reduced decoding when compared to hearing people. Dynamic Causal Modelling revealed that the ‘deaf’ motion-selective temporal region shows a specific increase of its functional interactions with hMT+/V5 and is now part of a large-scale visual motion selective network. In addition, we observed preferential responses to radial, compared to horizontal, visual motion in the ‘deaf’ right superior temporal cortex region that also show preferential response to approaching/receding sounds in the hearing brain. Overall, our results suggest that the early experience of auditory deprivation interacts with intrinsic constraints and triggers a large-scale reallocation of computational load between auditory and visual brain regions that typically support the multisensory processing of motion information.

**Highlights:**

- Auditory motion-sensitive regions respond to visual motion in the deaf
- Reorganized auditory cortex can discriminate between visual motion trajectories
- Part of the deaf auditory cortex shows preference for in-depth visual motion
- Deafness might lead to computational reallocation between auditory/visual regions.

## 1. Introduction

The study of the impact of sensory deprivation is a compelling model to investigate brain plasticity and the role experience plays in shaping the functional organization of the brain.

One fundamental discovery in the field of neuroscience has been that cortical regions deprived of their predisposed sensory input reorganizes by enhancing their response to the remaining senses (Bavelier and Neville, 2002). In the deaf brain, visual and somatosensory stimulation elicit enhanced activity in regions of the temporal cortex that are chiefly engaged in auditory processing in the hearing brain (Auer et al., 2007; Finney et al., 2001). However, what are the mechanisms governing the expression of such cross-modal plasticity remain debated (Cardin et al., 2020).

It has been proposed that the expression of cross-modal plasticity in sensory deprived people might be constrained by the innate domain specialization of the region lacking its predisposed sensory input (Lomber *et al*., 2010; Collignon *et al*., 2011; Reich *et al*., 2011; Ricciardi *et al*., 2014; Mattioni *et al*., 2020). Accordingly, in people who lack auditory experience, the auditory temporal cortex would maintain a division of computational labour that is somewhat similar to the one observed in the hearing brain (Dormal and Collignon, 2011). For instance, regions of the auditory cortex that typically process auditory localization in hearing cats respond selectively to visual localization in cats that were born deaf (Lomber et al., 2010). Only a few studies have provided evidence of functionally specific reorganization in the temporal cortex of deaf humans. For instance, Bola and colleagues reported preferential visual recruitment during rhythm discrimination when compared to non-rhythmic discrimination in a large portion of temporal regions (Bola et al., 2017). In another recent study, we reported preferential recruitment for faces in a discrete portion of the right mid-superior temporal sulcus that specifically responds to human voice in hearing subjects (Benetti *et al*., 2017; see also Stropahl *et al*., 2015 for similar results in postlingually deaf individuals with cochlear implant).

Although cross-modal activation of auditory regions has been observed in response to visual motion in deaf humans (Fine et al., 2005; Finney et al., 2001; Shiell et al., 2014), several important questions remain unanswered. In our study we tested whether visual motion processing recruits discrete regions of the temporal cortex known to selectively respond to specific auditory motion trajectories in hearing individuals (Battal et al., 2019; Hall and Moore, 2003; Warren et al., 2002). In addition, we used multivariate pattern decoding analyses to assess whether different motion trajectories could be discriminated in the deprived auditory cortex of deaf people. Importantly, we examined whether the cross-modal recruitment of specific temporal regions in the deaf was associated with reorganization of the computational capabilities within hMT+/V5 complex, which is known to implement multivariate responses to distinct motion trajectories (Kamitani and Tong, 2010, 2006). In deaf individuals, we also examined whether visual-motion recruitment of the temporal cortex was associated with reorganization of the large-scale cortico-cortical interactions with hMT+/V5 and ‘higher-level’ regions of the inferior parietal sulcus. Finally, by including a control group of proficient hearing signers, we could tear apart whether those potential changes were driven by congenital deafness or by the use of sign language.

## 2. Materials and Methods

### 2.1 Participants

Thirteen early deaf (ED), 15 hearing individuals (hearing-nSL), and 14 hearing fluent users of Italian Sign Language (hearing-SL) participated in the present fMRI study. The three groups were matched for age, gender, handedness (Oldfield, 1971), and non-verbal IQ (Raven et al., 1998). No hearing participants had reported neurological history, and all had normal or corrected-to-normal vision. Information on hearing status, history of hearing loss, and use of hearing aids were collected in deaf participants through a structured questionnaire. Early-deaf participants either self-reported their hearing loss based on the last audiometric testing or provided audiometric reports (see Table S1). Information about Sign Language age of acquisition and frequency of use was documented in both the deaf and hearing-signers group, and no significant differences were observed between the two groups (Table 1). In addition, since visual motion sensitivity and lip-reading skills seem to be related in deaf people (Mohammed et al., 2005), individual silent word lip-reading abilities were assessed in each group with a test we developed *ad hoc* for this study (LipIT, the Lip-Reading in Italian Test; *in preparation*).

**Table 1.** Demographics, lip-reading performances and Italian Sign Language aspects of the 42 subjects participating in the fMRI experiment.

| **Participants**<br>**fMRI**<br>**Experiment** | **Hearing-nSL**<br>**(n =15)** | **Hearing-SL**<br>**(n=14)** | **Early Deaf**<br>**(n=13)** | **Statistics** |
| --- | --- | --- | --- | --- |
| **Mean Age**<br>**Years (s.d.)** | 30.73 (5.49) | 34.64(5.74) | 32.77(7.50) | F-test= 1.415<br>*P-value* = 0.255 |
| **Gender (M/F)** | 8/7 | 4/10 | 6/7 | $\chi^2$= 1.897<br>*P-value* = 0.387 |
| **Non-verbal IQ**<br>**estimate (s.d.)** | 123.1 (8.65) | 124.8 (5.61) | 119.7 (9.92) | *K-W test* = 0.38<br>*P-value* = 0.827 |
| **Hand**<br>**preference,**<br>**% (right/left**<br>**ratio and s.d.)** | 69.31 (49.25) | 62.16 (52.73) | 60.83(53.53) | *K-W test* = 0.42<br>*P-value* = 0.810 |
| **LIS Exposure**<br>**Years (s.d.)** | -- | 25.00(10.25) | 21.08(14.37) | *t-test*= -0.078<br>*P-value* =0.439 |
| **LIS Acquisition**<br>**Age in years**<br>**(s.d.)** | -- | 12.33(8.69) | 9.64(11.35) | M-W.U-test = 101<br>*P-value* =0.381 |
| **Lip Reading**<br>**Composite**<br>**Z-scores (s.d.)** | -0.07(0.84)** | 0.47(0.56) ° | 0.70(0.51) | F-test= 5.210<br>*P-value* = 0.010 |

\*\* Significant at $p = 0.011$ Bonferroni-corrected in Deaf versus Hearing Controls; °Trend for significant difference at $p = 0.082$ in Hearing-nSL versus Hearing-SL. Abbreviations: LIS, Italian Sign Language;-nSL, LIS non-users -SL, LIS users; K-W, Kruskalis-Wallis; M-W, Mann-Whitney; s.d., standard deviation.

The study was approved by the Committee for Research Ethics of the University of Trento; all participants gave informed consent in agreement with the ethical principles for medical research involving human subjects (Declaration of Helsinki, World Medical Association) and the Italian Law on individual privacy (D.l. 196/2003).

### 2.2 Stimuli and Experimental Protocol

#### 2.2.1 Auditory Motion Localizer fMRI scan

In order to identify temporal regions sensitive to auditory motion and select the regions of interest (ROIs) for the main visual motion experiment, an independent group of 15 hearing-nSL individuals completed an auditory motion localizer fMRI scan. This experiment consisted of three main pink noise sound stimuli: static sound, lateral motion, and radial motion. Apparent perception of sound motion in the horizontal plane was achieved by modulating the relative intensity (stereo) of pink noise between the left and right ear (Interaural Level Difference, ILD); in the radial motion condition, sounds (mono) either rose or decreased exponentially in intensity creating the percept of a sound moving toward or away from the participant. A block design was implemented in a single run in which participants were asked to perform an oddball task and press a button when a sound moved slower or lasted longer (i.e., moving and statistic sounds respectively; 1.8s in duration) than the others (1s in duration) in the same block. The single run consisted of 30 consecutive blocks (10 repetitions/category) separated by rest periods of 9 s with no randomisation of trajectory presentation (order was: lateral, radial, static). Each block included 18 consecutive auditory stimuli (i.e. no interstimulus interval). Subjects were asked to maintain their sight on a central fixation cross for the entire duration of the experiment. A detailed description of the stimuli and of the experimental procedure for the auditory motion localizer can be found in the Supplementary Material (Section 1.1; Fig.S1) and in Dormal et al. 2016.

#### 2.2.2 Visual Motion fMRI Experiment

We developed an original fMRI paradigm of visual motion perception using global motion (Random Dots Kinematics; RDK) stimuli. The visual stimulation and the experimental paradigm were developed using Python language in OpenSesame (Mathôt et al., 2012), version Gutsy Gibson 2.8.1 (http://osdoc.cogsci.nl/). Video S1 provides an example of the stimuli used in each of the three conditions as well as the structure of stimulus delivery.

Global motion stimuli consisted of black dots on a gray background arranged in two circular arrays simultaneously displayed in the left and right hemispaces (Figure 1.A, C). The presentation of visual moving stimuli in both the left and right peripheral visual field had two main advantages: 1) previous studies have shown that peripheral visual stimuli are more prone to lead to cross-modal recruitments of temporal regions (Scott et al., 2014); 2) they allow the participant to compare the speed of motion between the left and right hemispaces and therefore psychophysically control performance level on a motion task using a staircase method (see below).

**Figure 1.**
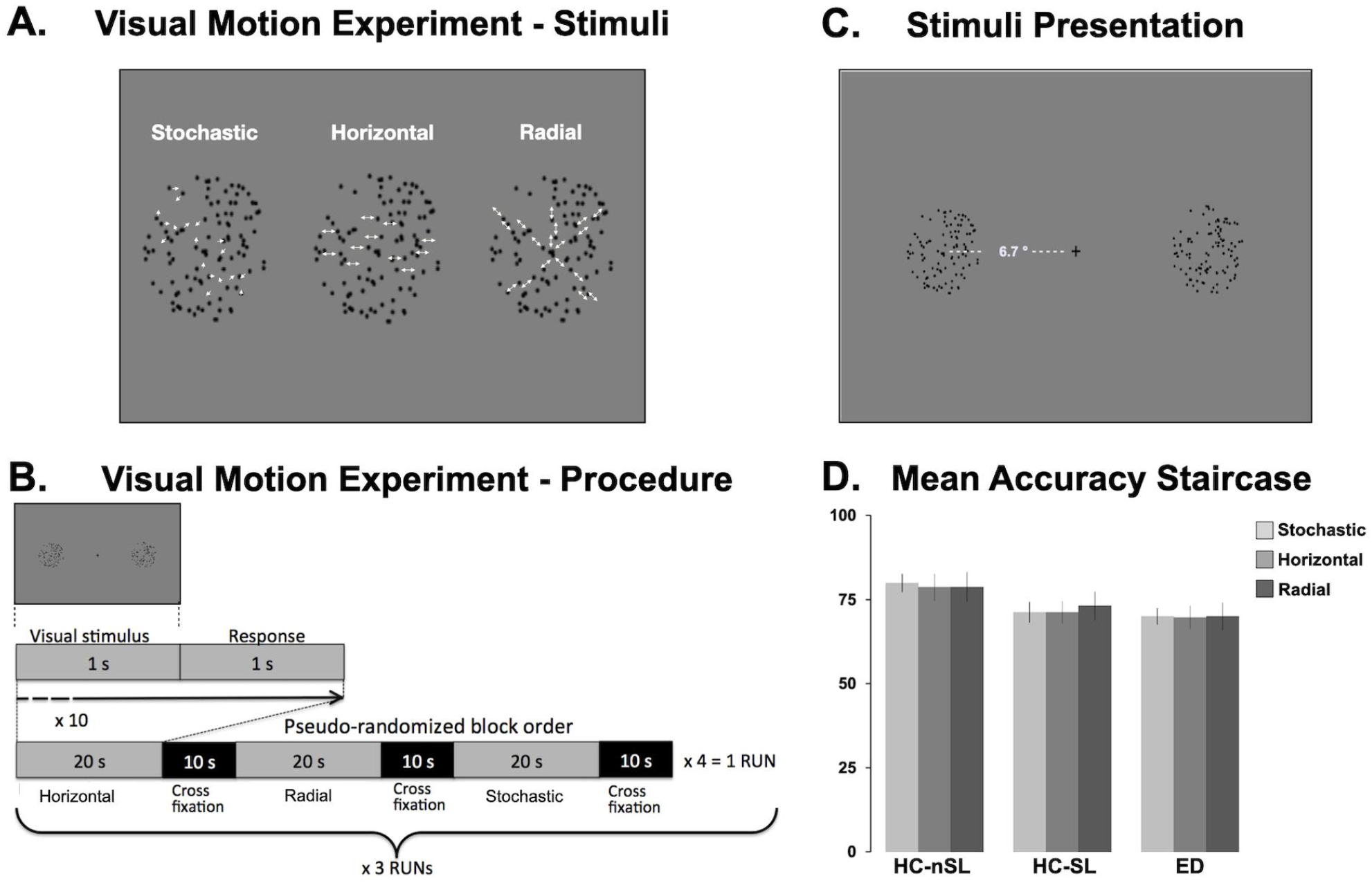
Illustration of the stimuli and procedure used in the visual motion experiment and mean group accuracies for the staircase procedure. **(A)** Schematic representation of the three experimental conditions: stochastic, horizontal and radial visual motion. Random Dots Kinematics were used as experimental stimuli. **(B)** Representations of a single trial and distribution of condition blocks within a run; condition blocks were pseudo-randomly ordered within a run to prevent contiguous repetition of the same block condition. **(C)** Examples of screen appearance during the presentation of the 2-alternative forced choice paradigm; dot arrays moved at different speeds based on the subject’s previous response. **(D)** Bar graphs show the mean accuracy across groups for the three main conditions: stochastic (light grey), horizontal (mid-grey) and radial (dark grey) motion. No significant differences between visual conditions or experimental groups were detected.

RDK stimuli were presented within visual frames of 1280 × 1024 pixels (about 17.81° × 14.29° of visual angle in the scanner) and with the following parameters: dot diameter = 0.07°, RDK diameter = 2.8°, RDK density = 16.24 dots/deg^2^. Three conditions were implemented: Stochastic Motion (SM), radial motion (RM), horizontal motion (HM). We included radial (dots alternatively approaching and receding) and horizontal (dots alternatively moving towards left and right) visual motion, since there is evidence that distinct neural populations respond to these motion trajectories both in audition (Stumpf et al., 1992; Toronchuk et al., 1992) and vision (Morrone et al., 2000; Saito et al., 1986; Tanaka and Saito, 1989), while stochastic motion was included to test the role motion coherence plays in eliciting brain responses (Tanaka and Saito, 1989; Morrone et al., 2000). To ensure that speed could not be determined from local motion detectors, the dots had a limited lifetime of 5 frames (∼83ms) and reappeared at a random position within the display area for a subsequent five frames lifetime (see Video S1). In the SM motion condition, dots disappeared and reappeared in a random position giving the sensation of stochastic flickering motion. In the RM the dots disappeared and reappeared coherently to create the sensation of approaching and receding motion. In the HM conditions, the dots disappeared and reappeared coherently to create the sensation movement in the left or right direction.

A block design was implemented in which participants were presented with a 2-Alternative Forced Choice paradigm and asked to indicate by button press which of the two dot arrays, simultaneously presented to the left and the right of the central fixation cross (0.7° × 0.7° degrees of visual angle in the scanner), appeared to move faster (Figure 1.A). Each dot array was centred at a visual eccentricity of 6.7° from central fixation (Figure 1.C) and had a radius of circular aperture of 1.4° in the scanner. Participants were asked to privilege accuracy over speed in responding but were however given a limited time for response since each trial was presented for 1s with an inter-stimulus interval of 1s. For each subject, three experimental runs were presented, each run containing four blocks per condition for a total of 12 experimental blocks in each run. Each condition-block consisted of 10 trials (therefore lasting 20s) and was alternated with an inter-block interval of 10s in duration consisting of a fixation cross. The condition-block order was pseudo-randomized, ensuring no consequent repetition of the same condition within the same run.

Crucially, a dynamic psychophysical staircase procedure was developed to control for task difficulty level across conditions and participants (Collignon et al., 2013, 2011), and cognitive aspects potentially impacted by early deafness (Brozinsky and Bavelier, 2004). This was achieved by adjusting the speed gap between each couple of dot arrays in each trial according to individual responses in the previous trial (1-step-down/correct response and 5-steps-up/wrong response; ∼70 % correct subject performance convergence, see Figure S2). For the initial trial of each of the first condition-block within a run, the speed difference was set to half the maximum difference range (4°-13°/s) for global motion since it maximally activates hMT+/V5 region within this range (Chawla et al., 1998).

### 2.3 Behavioural analysis

Task difficulty was tested by performing a mixed Analysis of Variance (ANOVA) in which motion trajectory was entered as within-subject repeated measure (3 conditions) and group as a between-subject factor (3 groups). Linear contrasts were then defined to test for main effects of group and condition as well as condition-by-group interaction (as implemented in JASP version 0.11.1, JASP Team, 2020).

### 2.4 fMRI data acquisition and analyses

#### 2.4.1 Data acquisition

Visual motion MR data was collected at the Center for Mind/Brain Sciences (CIMeC) using a Bruker BioSpin MedSpec 4T MR scanner and an 8-channel birdcage head coil. Images were collected with a gradient echo-planar sequence using a sequential ascending scheme (TR = 2200 ms; TE = 33ms; FA = 76°; slice thickness = 3 mm; FoV = 64×64 mm; 37 slices; 3 mm thickness; spacing between slices = 3.6 mm; slices = 792 for visual motion). In addition, a T1-weighted 3D MP-RAGE anatomical sequence was also acquired for coregistration purposes (TR = 2700 ms; TE = 4.18 ms; FA = 7°; slice thickness = 1mm; slices = 176). For the auditory motion fMRI localizer, functional MRI-series (acquired using a TRIO TIM system (Siemens, Erlangen, Germany) 3T MR scanner) were imported from a previous project of our group (Dormal et al., 2016). See a detailed description of data acquisition parameters in supplementary material (Section 1.2).

#### 2.4.2 Univariate analysis of fMRI data

Functional datasets from the visual and auditory experiments were pre-processed and analysed using SPM12 (http://www.fil.ion.ucl.ac.uk/spm12) and Matlab R2012b (The Matworks, Inc). In each functional dataset, the images were corrected for differences in slice acquisition timing, motion corrected (6 parameters affine transformation) and realigned to the mean image of the corresponding sequence. The individual T1 image was segmented in grey and white matter parcellations and the forward deformation field computed. Functional EPI images (3mm isotropic voxels) and the T1 image (1mm isotropic voxels) were normalized to the MNI space using the forward deformation field parameters and data resampled at 2mm isotropic with a 4^th^ degree B-spline interpolation. Finally, the functional images in each dataset were spatially smoothed with a Gaussian kernel of 6mm full width at half maximum (FWHM).

Data analysis was implemented within the general linear model (GLM) framework with a mixed effects model, which accounted for both fixed and random effects. For the single-subject analysis, we defined a design matrix including separate regressors for the conditions of interest (see below), plus realignment parameters to account for residual motion artefacts as well as outlier regressors; these regressors referred both to scans with large mean displacement and/or weaker or stronger globals (Carling, 2000; Siegel et al., 2014). The regressors of interest were defined by convolving boxcars functions representing the onset and offset of stimulation blocks in each experiment by the canonical hemodynamic response function as implemented in SPM. Each design matrix also included a filter at 168s and auto-correlation, which was modelled using an auto-regressive matrix of order 1.

**For the auditory localizer**, three predictors were modelled and linear contrasts defined to test: the main effect of global motion processing ([lateral + radial > static]) and the main effect of motion-direction processing ([radial *versus* lateral]). These contrast images were then further spatially smoothed by a Gaussian kernel of 6mm FWHM prior to group-level analyses, where they were inputted in a series of corresponding one-sample *t*-tests within the independent hearing group. Statistical maps were corrected for multiple comparisons at the whole-brain level using cluster-level p < 0.05 family-wise error (FWE) correction and a defining voxel-level threshold of p < 0.001. Only clusters with a minimum size of 20 contiguous voxels (160mm3) were selected (Flandin and Friston, 2016). The regional group-maxima peak-coordinates for each contrast were taken as the centre of the three ROIs used in the analysis of the visual-motion experiment: the A-motion-STC, A-radial-STC, and A-lateral-STC (Figure 2; see Results sections for a detailed description of auditory ROIs and Table S2 for ROI coordinates).

**Figure 2.**
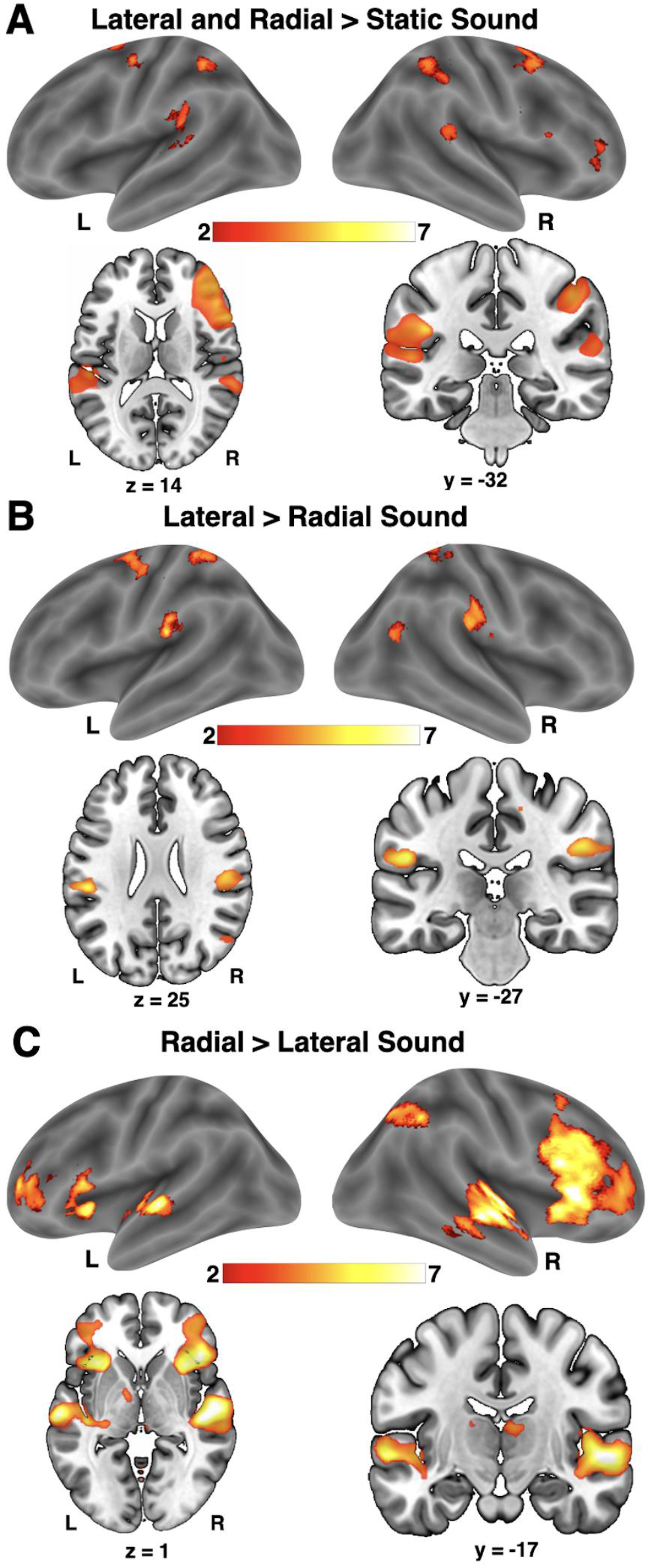
Results of the whole brain univariate analysis for the auditory motion experiment in hearing-nSL individuals. Regional responses are superimposed on renders (top panels) and axial/coronal slices (bottom panels) of the MNI-ICBM 12 brain template (SurfIce, https://www.nitrc.org/projects/surfice/) reported for visualization purposes at *P* < 0.005-uncorrected, cluster size k = 20. T-values are scaled according to the colour-bar. L, Left; R, Right

**For the visual task**, condition predictors were modelled at the subject-level and linear contrasts defined to test: the main effect of each coherent radial and horizontal motion condition ([RM > SM], [HM > SM]), the main effect of coherent motion ([RM+HM] > SM) and the main effect of visual motion ([RM+HM+SM > static fixation-cross]). These contrast images were further spatially smoothed by a 6mm FWHM Gaussian kernel prior to enter group-level analyses. At the group-level, we performed a series of one-sample *t*-tests to examine main effects of each coherent motion trajectory within individual groups. Subsequently, two one-way Analyses of Variance (ANOVAs) were implemented to detect group differences for coherent motion, and all visual motion conditions in [early deaf > hearing-nSL] conjunction [early deaf > hearing-SL]. In addition, we looked at the Trajectory-by-Group interaction effect by implementing a mixed Analysis of Variance (factorial design) with global motion direction as within-subject factor and the three groups as between-subject factor ([RM *vs*. HM] in [early deaf *vs*. hearing-nSL] conjunction [RM *vs*. HM] in [early deaf *vs*. hearing-SL]).

In our main ROIs analysis, group differences were tested at a statistical inference of *P* < 0.05 FWE voxel-corrected over a small spherical volume located at the peak coordinates for the relevant ROI extracted from the auditory motion localizer (i.e. A-motion-STC, A-radial-STC or A-lateral-STC, see Fig. 2). A radius of 9 mm for each sphere was applied in order to account for inter-individual variability in functional localization while avoiding ROIs overlapping and ensuring a sufficient level of anatomical specificity. Significant effects were further characterized by extracting mean weight parameters for the relevant conditions and statistically assessing effect size and relative predictive performance of competing hypotheses (JASP Version 0.11.1, JASP Team, 2019; Wagenmakers et al., 2018b, 2018a).

Additionally, to assess the spatial selectivity of the observed effects and characterize potential differences between groups occurring outside the selected ROIs, we computed a more explorative whole-brain analysis for which statistical inference was performed at P < 0.001 (uncorrected for multiple comparison) with a cluster size of a minimum of 20 contiguous voxels (160mm^3^).

#### 2.4.3 Multivariate pattern analysis

Since the univariate analysis revealed recruitment of the right STC in response to global visual motion but no selectivity to the coherence of visual motion (i.e. [HM+RM>SM]) in deaf individuals, we used multivariate pattern analysis in order to explore whether this region contains reliable distributed information about motion planes and coherence. We also aimed at examining whether cross-modal recruitment of the right auditory region in the deaf was associated with reorganization of the computational capabilities within the ipsilateral regions typically dedicated to visual motion processing (i.e. right hMT+/V5; see Dormal et al. for similar reasoning with blind participants). The pre-processing steps were identical to the pipeline applied to the univariate analysis, except no smoothing was applied to the functional data. A GLM model was implemented including a separate regressor for each block in each run of the visual experiment (i.e. 3 experimental conditions × 4 repetitions × 3 runs = 36 independent regressors, with the exception of one participant who underwent only 2 runs = 24 independent regressors). Finally, for each block in each run, t-maps were estimated for each condition using the contrast [motion condition > baseline]. The t-maps were used for further multivariate pattern analysis (MVPA).

The hMT+/V5 ROI was defined as a 9 mm sphere around the group-maxima for visual motion selectivity [RM + HM+SM > fixation cross] in the conjunction analysis across the three groups (MNI coordinates: [48 −67 5]). The A-motion-STC ROI was defined as a 9 mm sphere around the group-maxima of the contrast in [Early Deaf > Hearing-nSL and Hearing-SL] for [visual motion > fixation-cross] masked by the contrast for [auditory motion > static sound] in hearing individuals (MNI coordinates:[60 −34 11]). This masking procedure ensures that the region under investigation with multivariate analysis is 1) showing enhanced cross-modal response in deaf participants when compared to hearing people and 2) is a brain region showing auditory motion selectivity in the hearing brain. The choice of spherical ROIs was to equate the number of features/voxels across subjects and groups to eliminate any bias in the decoding accuracy that could originate from having more features in one of the groups. The decoding analysis was performed using CoSMoMVPA (http://www.cosmomvpa.org; Oosterhof et al., 2016) implemented in Matlab. Classification was performed using a linear discriminant analysis (LDA) classifier as it was demonstrated that linear classifiers perform better than other non-linear or correlation based methods (Misaki et al., 2010). The LDA classifier was trained and tested to discriminate the 3 motion conditions in each subject; the training for discrimination was performed on two runs and the testing on the left-out run (leave-one run out). The previous step was repeated 3 times (N-fold cross-validation) with the classifier tested on a different run in each cross-validation and single classification accuracy obtained by averaging the classification accuracy from all the cross-validation folds. The resulting three-class classification accuracy was further analysed as follows. First, a one-sided one-sample t-test was performed to assess whether the classification accuracy was significantly above chance level (33.33%) in each ROI and for each Group. Second, a mixed-effects repeated measure ANOVA was implemented in which decoding accuracy was entered as the dependent measure, Region (ROI) as a within-subject independent factor and Group as a between-subject independent factor. Significant main effects and interactions were subsequently characterised by implementing a series of post-hoc t-tests. FDR correction (Benjamini and Yekutieli, 2001) was applied to the multiple comparisons accordingly. Finally, in order to further evaluate the reliability of the parametric assessment and the significance of the estimated decoding accuracies, we also implemented an additional non-parametric assessment and applied a binomial formula to the data as previously recommended (Combrisson and Jerbi, 2015). Full details of these additional analyses and of their results can be found in the Supplementary material (Sections 1.3 and 2.4).

#### 2.4.4 Effective connectivity analysis

The aim of dynamic causal modelling (DCM; Friston *et al*., 2003) is to estimate, and make inferences about, the influence that one neural system exerts over another and how this is affected by the experimental context. In dynamic causal modelling, a reasonably realistic but simple neuronal model of interacting neural regions is constructed. Dynamic causal modelling uses a previously validated biophysical model of fMRI measurements to predict the underlying neuronal activity from the observed hemodynamic response; the estimated underlying neural responses are then used to derive the connectivity parameters (Friston, 2011; Stephan, 2006). In combination with Bayesian model selection (BMS) and Parametric Empirical Bayes (PEB; Friston *et al*., 2016), DCM can be used for the comparison of competing mechanistic hypotheses of brain connectivity, represented by different network models and different model families (Penny et al., 2010, 2004), as well as for inference on estimated group-level connectivity parameters (Zeidman et al., 2019). Here, we adopt this approach to answer three fundamental questions in early deaf individuals. At the parameter-level and across deaf individuals, we characterized a common effective connectivity profile associated with visual motion processing within a network of regions including, the right A-motion-STC, the right hMT+/V5, and the right intraparietal sulcus (IPS). At the model-level, we first examined whether, in deaf individuals, visual motion inputs reach the reorganized A-motion-STC directly from subcortical structures or through connections with motion-selective hMT+/V5. Finally, we tested whether, despite the absence of different univariate responses across motion conditions in A-motion-STC, coherent visual motion processing modulates the strength of coupling between right hMT+/V5 and A-motion-STC in the audio-visual motion network of deaf individuals.

DCM models can be used for investigating only brain responses that present a relation to the experimental design and can be observed in each individual included in the investigation (Stephan et al., 2010). Because no temporal activation was detected for visual motion processing in hearing controls, they were not included in the DCM analysis. To the purpose described above, we first defined a network of regions including the right hMT+/V5, the right A-motion-STC and the right IPS, based on the observed regional responses to visual motion in the deaf group (see figure 3B, supplementary material and Table S2 for details on ROIs definition).

**Figure 3.**
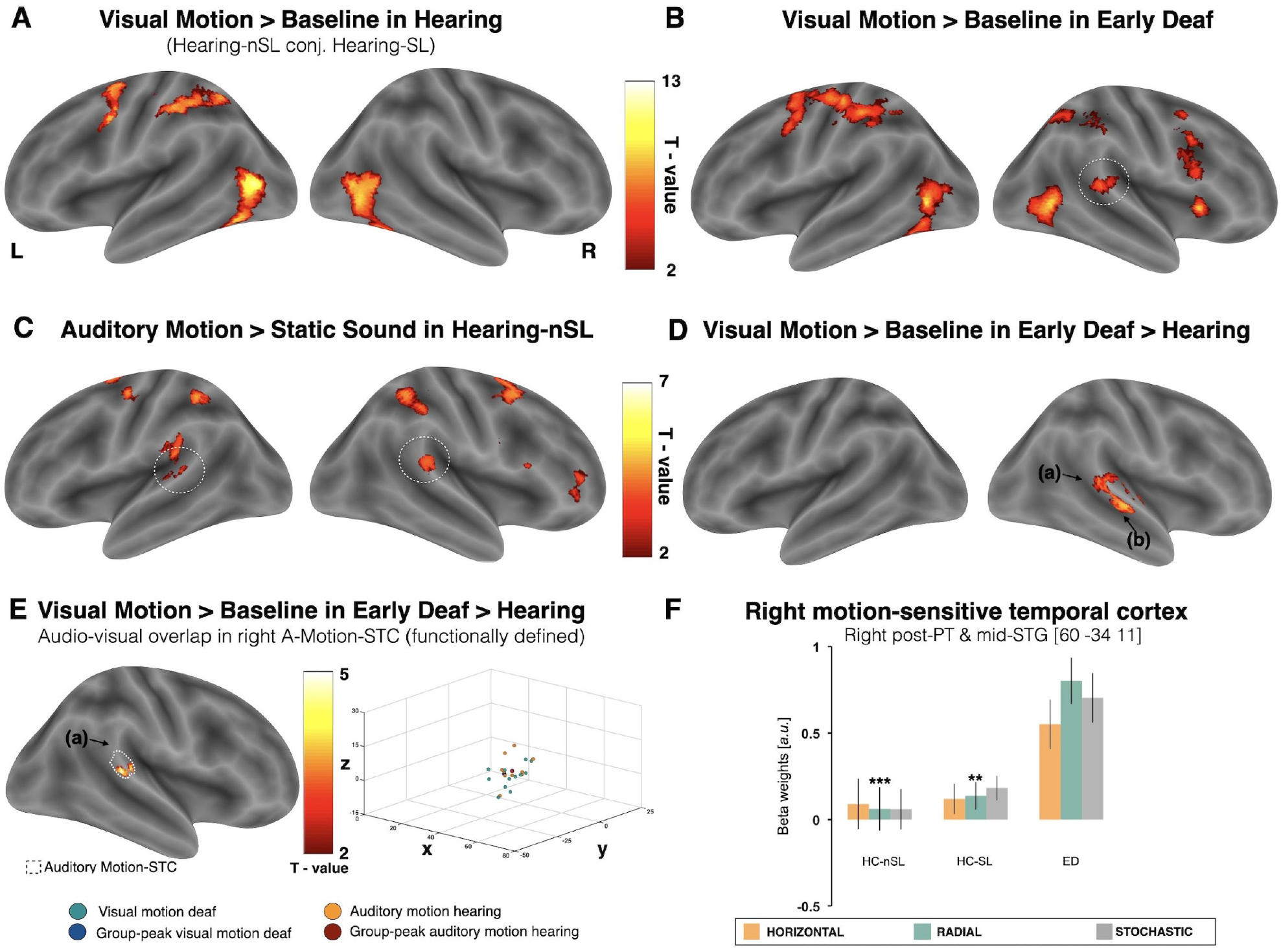
Visual motion activates the temporal auditory cortex in early deaf individuals. **(A) and (B)** Global visual motion activity in the hearing (merged) and deaf groups (displayed at *P*_unc._ < 0.001, cluster size > 20 for visualization purposes). **(C)** Auditory motion activity (conjunction of looming and lateral moving sound *vs*. static sound) in an independent group of hearing-nSL individuals, the auditory motion-sensitive temporal cortex is circled in dotted-line (displayed at *P*_unc._ < 0.005, for visualization purposes). **(D)** Group-by-Task interaction, visual motion compared to visual baseline showed significant activation only in the right temporal cortex of early deaf compared to hearing controls (displayed at *P* < 0.05 FWE cluster-corrected, cluster size > 20); the crossmodal recruitment encompassed lateral and posterior *planum temporale* (a) and mid-superior temporal gyrus and sulcus (b). **(E - left)** Visual motion sensitivity within the right deaf A-motion-STC (*P* <0.05 FWE over SVC); outlined in black dotted-line is the auditory motion-STC as functionally defined in hearing controls for [auditory motion > static]. (**E – right)** 3D scatterplot depicting the overlap of individual activation peaks in right STC for visual motion in early deaf subjects (turquoise circles) and auditory motion in hearing individuals (orange circles); dark blue and dark red circles represent the group-maxima of each group. **(F)** For visual purpose only, mean response estimates (a.u. +/- SEM) during visual motion are reported for hearing and deaf groups in temporal region showing the strongest motion-sensitive cross-modal recruitment; Bonferroni-corrected *P-values* for ED>HC-nSL and ED>HC-SL: ** *p <* 0.01 and *** *p <* 0.001. Abbreviations: STC, superior temporal cortex; PT, *planum temporale*; STG, superior temporal gyrus; HC-nSL, hearing non-signers; HC-SL, hearing signers; ED, early deaf; L, left; R = Right; A, anterior; P, posterior.

At the model and family levels, we defined a set of two family-features (see figure 6A): (a) models with visual input to right hMT+/V5 only or both right hMT+/V5 and A-motion-STC, and (b) models with a modulatory effect of coherent motion on hMT+/V5-STC coupling or without it. Subsequently, we operationalized our model space based on the factorial combination of family features with each model falling within four possible families (see figure 6A and supplementary material). In addition, for each model a version with forward modulation only and one with both forward/backward modulation between regions were included. This process resulted in a set of 18 DCM models that were fitted to the fMRI data of the 13 deaf individuals for a total of 234 fitted DCMs and corresponding log-evidence and posterior parameter estimates. Subsequently, a two-folded BMS (see supplementary material, Section 1.5) was implemented to identify the winning family and the model that better explained the regional responses observed in deaf individuals.

At the parameter level, since we had no *a-priori* hypothesis on which connections prominently characterised the connectivity profiles associated with visual motion processing in the deaf group, we performed an automated search over estimated group-level connectivity parameters by using PEB in combination with Bayesian Model Reduction (BMR) and Bayesian Model Average (BMA). The group-level estimated connectivity parameters were then thresholded for significance based on the free energy criterion at a posterior probability <0.95% (Friston et al., 2007), resulting in a profile of excitatory and inhibitory connectivity changes that significantly contributed to the observed cross-modal responses in the auditory motion-STC of deaf individuals (see supplementary material for a detailed description of PEB implementation in this study; Section 1.6).

## 3. Results

### 3.1. Behavioral Results

#### Visual Motion Experiment

The mixed ANOVA on group mean accuracy values revealed that there were no significant main effects either of motion conditions/trajectories (F=0.512, *P*=0.601) or groups (F=2.021, *P*=0.146) and no interaction between those two factors (F=0.687, *P*=0.603), supporting that our psychophysical staircase procedure inside the scanner allowed for a comparable performance amongst the experimental groups (Figure 1.D).

#### Lip-reading Test

A one-way ANCOVA was implemented with a composite measure of accuracy on single words and sentences (in z-scores) as dependent variable, groups as between-subject factor and age as a covariate of no interest. A significant main effect of group was found (F = 5.210, *P* = 0.010) and, subsequently, a series of *post-hoc* two-sample *t*-test revealed that deaf participants performed significantly better than hearing controls (t = 3.101, *P* = 0.011) but not than hearing-SL (t = 0.779, *P* = 0.441) participants. A weak trend for significant difference was also observed between the two hearing groups (t = −2.297; *P* = 0.082). However, no significant correlation was observed between individual composite lip-reading measures and individual mean accuracy values on the visual motion experiment (R = −0.027, P = 0.866).

### 3.2. fMRI results – Univariate Analyses of Auditory Motion data

Consistent with previous work (Hall and Moore, 2003; Pavani et al., 2002; Warren et al., 2002), the independent auditory motion localizer showed selective activity for both lateral and radial motion along the anterior-posterior axis of the *planum temporale* bilaterally extending ventrally to the superior bank of the superior temporal sulcus (Figure 2). The selective response to lateral auditory motion was observed bilaterally in the most posterior portion of the STC extending dorsally to the supramarginal gyrus (MNI coordinates: right [48 −28 28], left [−52 −26 22]; Fig. 2B) while the selective response to radial auditory motion peaked more anteriorly in the STC extending ventrally to the mid superior temporal sulcus (STS; MNI coordinates: right [56 −17 −2], left [−56 −24 −4]; Fig. 2C). Hereafter, these regions will be referred to as A-lateral-STC and A-lateral-STC respectively. In addition, a sub-region responding to both lateral and radial motion was functionally localized in the mid-posterior and lateral portion of the STC, bilaterally, and will, hereafter, be referred to as the A-motion-STC (MNI coordinates: right [64 −36 14], left [−56 −28 8]; Fig.2A). These regions correspond to the auditory regions used for the ROIs analysis of the main visual motion experiment.

### 3.3. fMRI results – Univariate Analyses of Visual Motion data

#### Regions for visual motion processing common to all groups

Consistent with previous studies (Braddick et al., 2001; Rees et al., 2000; Watson et al., 1993; Zeki et al., 1991) comparing visual motion with static fixation-cross engaged a network of regions including the bilateral hMT+/V5 and intraparietal sulcus in both hearing and deaf individuals (Figure 3A and 3B; Table 2). Since no significant differences were observed between the two hearing groups, we pooled hearing-nSL and hearing-SL individuals, hereafter referred to as ‘hearing group’, to exclusively investigate the specific effects of sensory deprivation on cross-modal reorganization of auditory regions (Cardin et al., 2013) and reduce multiple comparisons issues in our statistics.

**Table 2.**
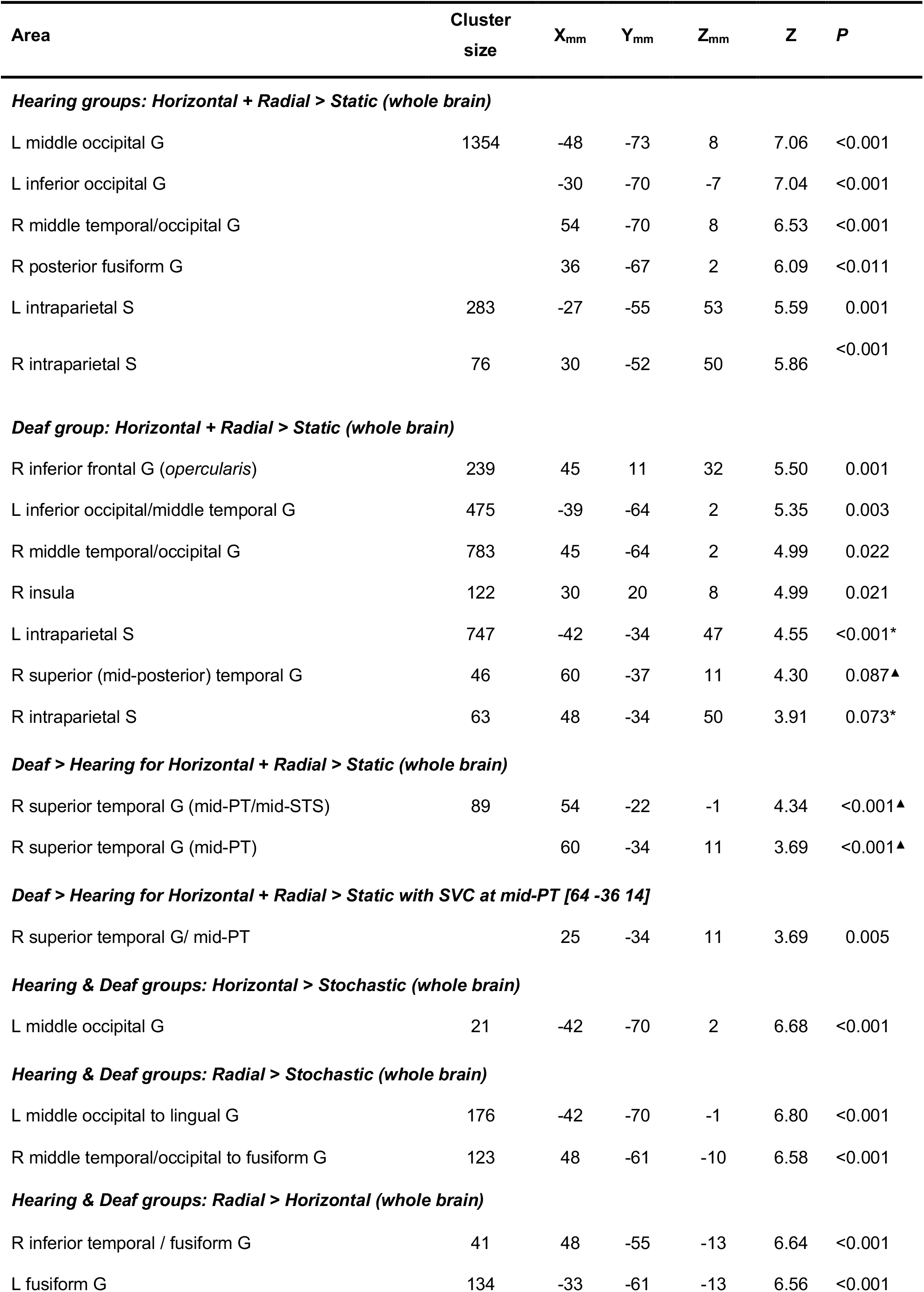
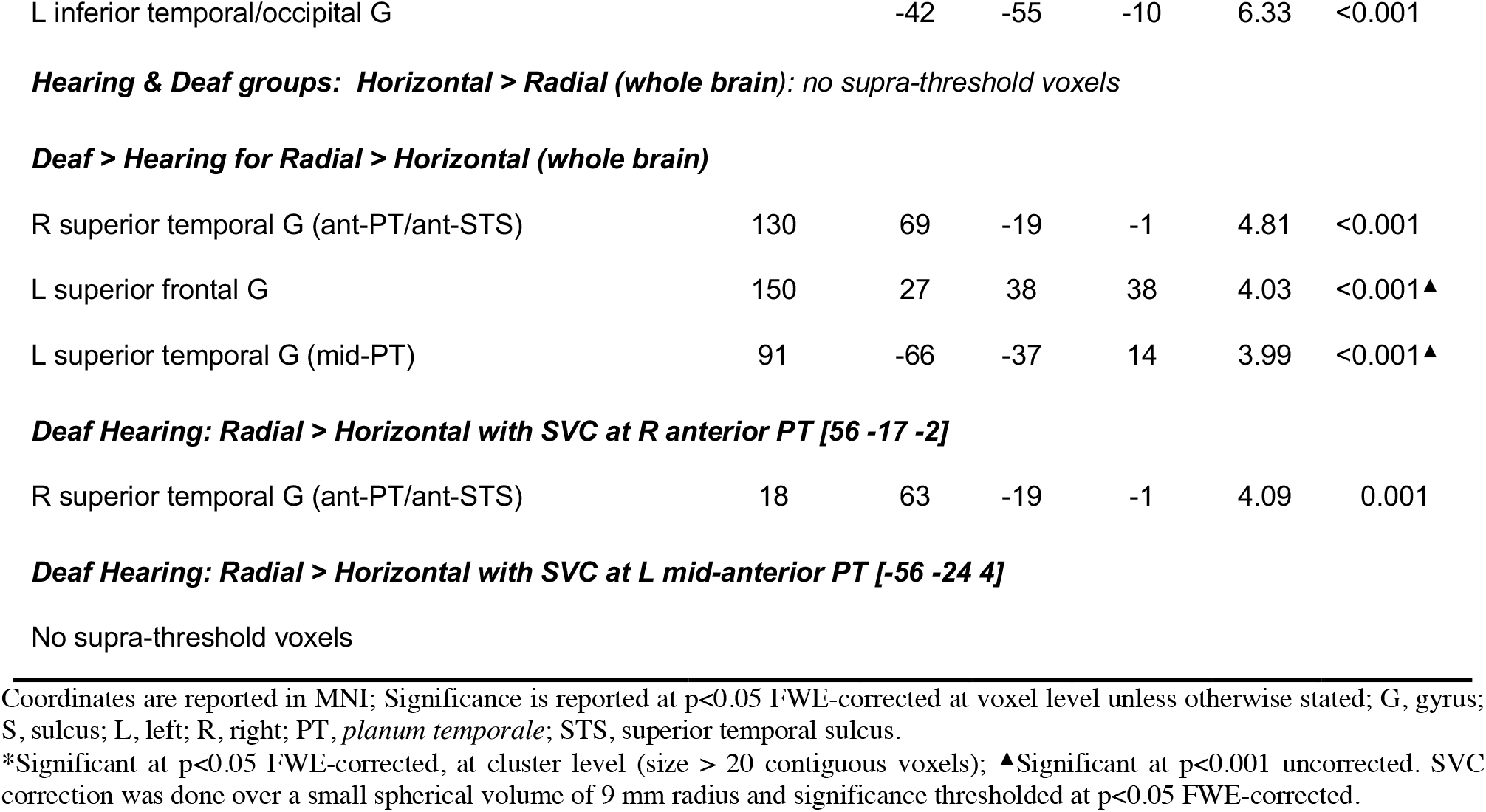
Results of the univariate fMRI analyses for the visual motion experiment. Regional responses for the main effect of visual motion and specific visual motion conditions are reported in hearing and deaf groups along with group differences.

#### Cross-modal recruitment of the temporal region during motion processing in deaf individuals

In deaf participants in addition to the occipital and parietal clusters observed in all groups, a cluster of visual motion response was detected in the STC (Circled in Figure 3B) which overlapped with the temporal region responding to auditory motion in the hearing group (Circled in Figure 3C and statistically isolated in figure 3E). Repeating this comparison across the whole brain confirmed a cross-modal recruitment restricted to the right STC in the ventral PT and extending anteriorly to the superior bank of the STS (MNI coordinates: right [57 −22 1] to [60 −34 11]) of early deaf compared to hearing individuals (Figure 3D, Table 2). This anterior effect, however, did not survive FWE-cluster correction at the whole brain level. To further describe the visual motion response observed in the right temporal cortex, we extracted individual measures of mean estimated activity (beta weights) in response to global motion (i.e. HM, RM and SM) from the right A-motion-STC as independently localized in the hearing groups.

In these regions, an analysis of variance revealed a main effect of group (F = 16.18, P < 0.001, η_p_^2^ = 0.310; Figure 3F), confirming increased motion selective response in this region of deaf compared with both the hearing-nSL (*t* = 3.425, P < 0.002; Cohen’s d = 1.298) and hearing-SL individuals (*t* = 3.771, P < 0.001, Cohen’s d = 1.452). The estimation of the Bayes Factor (BF) additionally revealed that the main effect of group had a BF_10_ = 44, suggesting strong evidence in support of cross-modal recruitment of the ‘deaf’ A-motion-STC for visual motion (see Section 2.3.1 in Supplementary Material for further details on the Bayesian analysis).

#### Responses in the A-radial-STC during radial visual motion processing in deaf individuals

In order to further test for motion-trajectory selectivity, we contrasted RM and HM conditions against each other. No selective recruitment for horizontal motion was observed within the lateral-STC in early deaf compared to hearing individuals. In contrast, we observed a selective response in ED for radial motion within the right A-radial-STC ROIs (Figure 4A-B, Table 2). Extending the comparison to whole-brain analyses confirmed the observations made at the ROI level but revealed additional foci of sub-threshold cross-modal activations in the left posterior STC and in the superior frontal sulcus (Figure 4A, table 2). The inspection of individual peak functional responses to visual radial motion in deaf individuals and auditory radial motion in hearing controls revealed a high degree of spatial overlapping (Figure 4C). To further describe the effect size of selective responses to radial motion in the mid-STC, we extracted individual measures of mean estimated activity (beta weights) in response to radial and horizontal motion (i.e. RM and HM) from the right A-radial-STC as independently localized in the hearing groups. In these regions, a mixed-effect repeated-measures analysis of variance revealed a significant [Group-by-Condition] interaction (F = 15.29, P < 0.001, η_p_^2^ = 0.440; Figure 4D), confirming increased responses to radial (F = 9.841, p < 0.001, η_p_^2^ = 0.327), but not horizontal (F =1.873, p = 0.167) motion in this region of deaf compared to hearing individuals. A Bayesian mixed-effect ANOVA additionally revealed a BF_10_ = 1174 for the interaction term (Group-by-Motion) indicating very strong evidence in favour of a selective response to radial, as opposed to horizontal, visual motion in the ‘deaf’ A-radial-STC (for further details see Section 2.3.2 in Supplementary Material).

**Figure 4.**
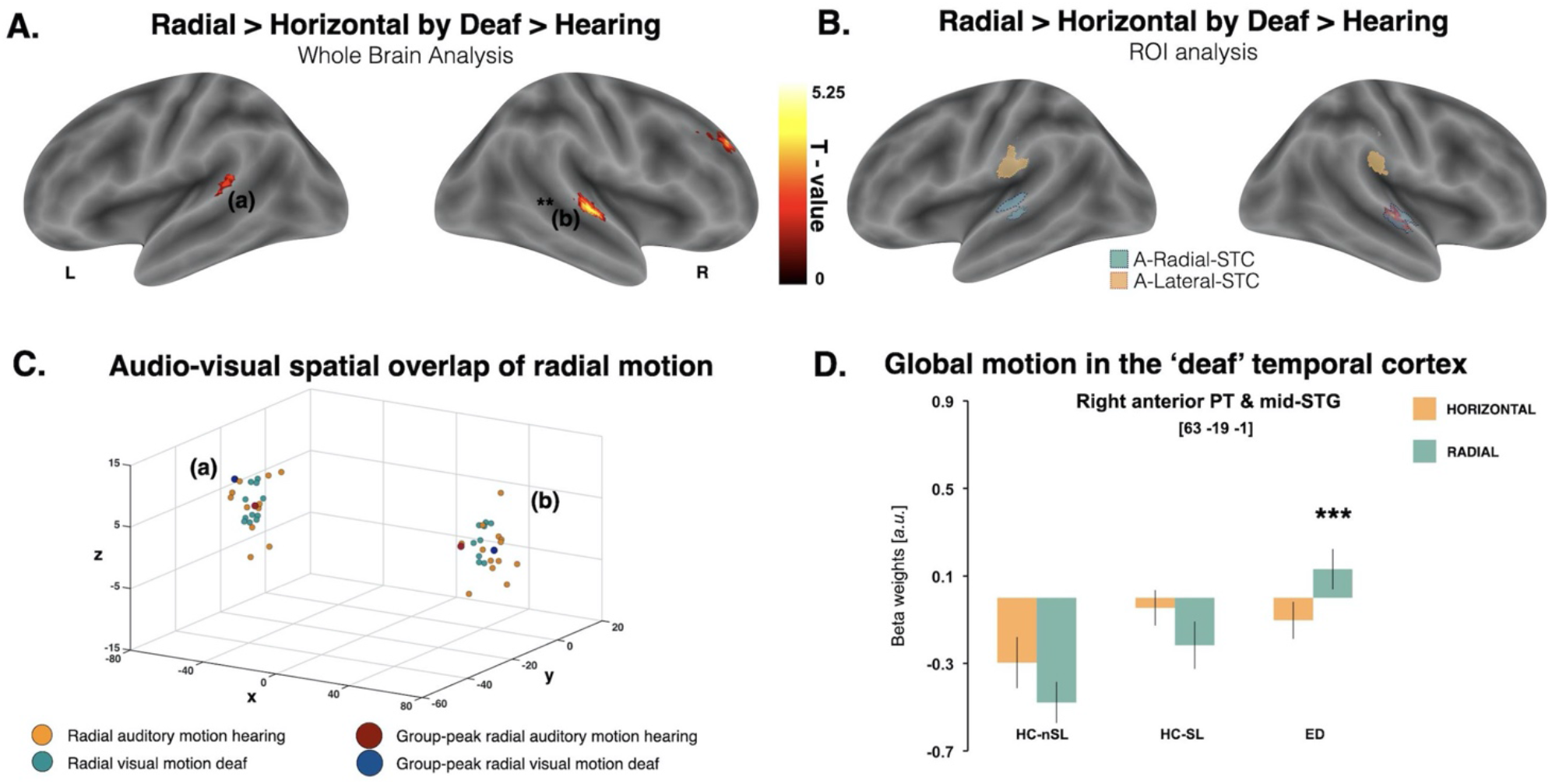
Cross-modal functional preference for radial visual motion. **A**. Whole-brain radial visual motion responses in the left posterior PT (a) and right anterior-PT/mid-STG (b) of deaf individuals (*P* <0.001 uncorrected for visualization purposes; ** significant at P<0.05 FWE-corrected); **B**. Cross-modal activations to radial visual motion in the right A-looming-STC (*P <* 0.05 FWE-corrected over small volume, radius 9mm); outlined are the auditory looming-STC (in turquoise) and the A-lateral-STC (in orange) ROIs. **C**. 3D scatterplot depicting individual activation peaks in PT/STG for radial visual motion in early deaf subjects (turquoise circles) and radial auditory motion in hearing individuals (orange circles); dark blue squares and dark orange circles represent the group-maxima of each group. **D**. For completeness, mean response estimates (a.u. +/-SEM) during radial visual motion are reported for hearing and deaf groups in the right temporal region showing cross-modal responses; Bonferroni-corrected *P-values* for radial > horizontal motion in ED>HC-nSL and ED>HC-SL: *** *p <* 0.001. Abbreviations: STC, superior temporal cortex; PT, *planum temporale*; STG, superior temporal gyrus; HC-nSL, hearing non-signers; HC-SL, hearing signers; ED, early deaf; L, left; R = Right; A, anterior; P, posterior.

### 3.5. fMRI results – Multivariate pattern analysis and motion decoding

Since an enhanced cross-modal response was observed in ED to all motion conditions *versus* static fixation-cross within the right deaf motion-STC, we relied on multivariate pattern analyses to further investigate the presence of visual information for motion direction in deaf compared to hearing individuals. Moreover, we examined whether such reorganization impacts on the computational capabilities of the right hMT+/V5 area, one of the most prominent regions coding for those different motion conditions.

In this analysis, the hearing-nSL and hearing-SL groups were pooled in one hearing group since no significant differences were observed between them. We observed a significant Group-by-Region interaction (F = 5.61, *P* = 0.020; Figure 5). This interaction was driven by higher decoding accuracy in early deaf than in hearing individuals in A-motion-STC, while the opposite profile was observed in hMT+/V5. Post-hoc comparisons showed that in A-motion-STC decoding accuracy of motion direction was above chance level only in the deaf group (deaf: T = 3.126, *P =* 0.010; hearing: T = 0.998, *P* = 0.327; Figure 5 bottom-left) and was significantly higher than hearing people (T = 2.134, *P* = 0.039). However, in hMT+/V5, decoding accuracy of motion direction was significantly above chance in both groups (deaf: T = 2.985, *P =* 0.012; hearing: T = 0.868, *P* < 0.001; Figure 5 bottom-right) and no significant difference was observed between the two groups (T = −1.208, *P =* 0.234).

**Figure 5.**
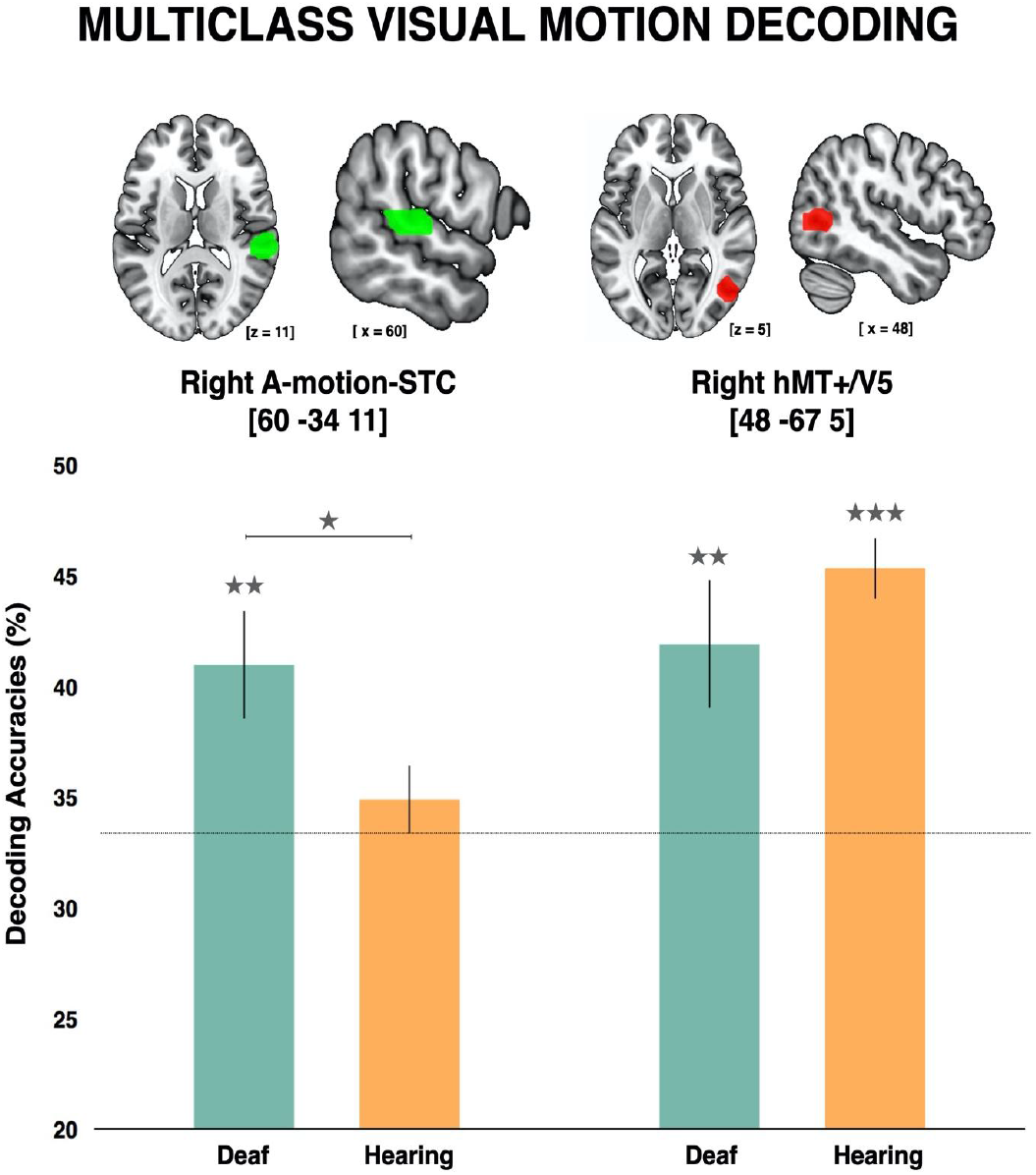
Multiclass visual motion decoding within the right A-motion-STC and right hMT+/V5 across the three groups. Bar graphs show the mean decoding accuracy (%*±* SEM) of the 3 motion visual conditions (FD, GHM, GHM,) in each group for the two regions, which are shown superimposed on an MNI-ICBM 12 brain template on the top. Uncorrected p-values: *p 0.05, **p 0.01, ***p 0.001 testing differences against chance level (33.33%; in dotted lines) and between groups.

### 3.4. fMRI results – Dynamic Causal Modelling

#### Model level

The two-folded random-effects family comparisons, addressing the target region/s of visual motion input and the modulatory effects of visual coherent motion on network connections, revealed that models including visual input to both the hMT+/V5 and the A-motion-STC as well as modulation of coupling strength between these two regions during coherent motion best explained the cross-modal visual responses observed in the reorganized A-motion-STC of deaf individuals (Family 4; Fig. 6B). Within the selected family, the random-effects BMS showed that the model including modulatory effects of coherent motion on reciprocal connections between each region in the motion network (Model 18) had the highest probability of evidence (exceedance probability = 0.754; Fig. 6C).

**Figure 6.**
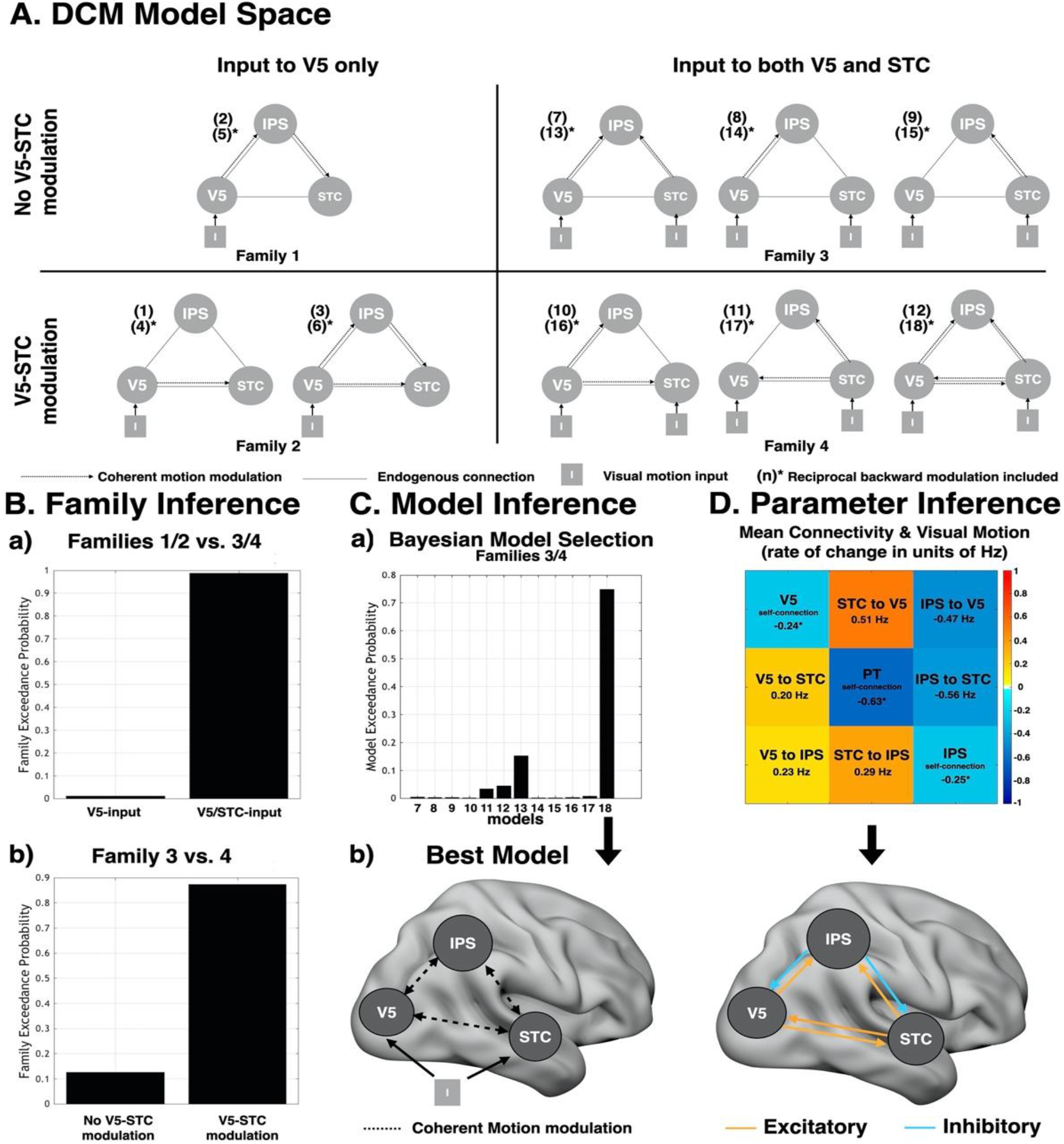
Dynamic Causal Modelling analyses. **(A)** The 18 DCMs used for Bayesian comparison in this study and the subdivision of the model space in four alternative families depending on the factorial combination of visual input target regions and coherent motion modulatory effects. Models 4-6 and 13-18 included forward/backward connections and are not depicted here to avoid visual redundancy. **(B)** Two-folded family-wise Bayesian selection, showing that visual motion input to both V5/STC and coherent motion modulatory effects on V5-STC connections best fit the regional responses observed in deaf individuals. **(C)** Random effects Bayesian model selection, revealing that modulation of both forward/backward connectivity between each region in the reorganized motion network best explained the observed data in the deaf group. **(D) Top**. PEB connectivity matrix reporting the self- and extrinsic connections significantly contributing to visual motion processing across deaf individuals. Excitatory or inhibitory effects are quantified in rates of change (in units of Hz) in the target region. Self-connections are quantified in log of scaling parameters that multiply up or down the default value of −0.5Hz; negative values indicate less self-inhibition during visual motion processing. A schematic representation of the extrinsic connectivity profile is also depicted **(D) Bottom**. Abbreviations: V5, hMT+/V5 region; STC, A-motion-STC; IPS, intraparietal sulcus; I, visual motion input.

#### Parameter level

The automated PEB search over group-level connectivity estimates revealed a connectivity profile, common to all deaf individuals and averaged across motion trajectories, in which excitatory connections between visual and auditory motion-selective regions, as well as excitatory/inhibitory connections between these regions and the right IPS, significantly contributed to the cross-modal responses to visual motion observed in the right A-motion-STC (Figure 6D). Interestingly, the strength of the excitatory influence that the reorganized A-motion-STC exerted over the hMT/V5 almost doubled its reciprocal from the hMT+/V5 (Fig.6D-top).

## 4. Discussion

### 4.1. Functional responses for visual motion in right STC of deaf subjects

Consistently with previous investigations (Fine et al., 2005; Finney et al., 2001; Sadato et al., 2005; Shiell et al., 2014), the present work confirms the recruitment of the superior temporal cortex in response to visual motion processing in early deaf individuals. Our study adds on these previous reports by showing that cross-modal visual motion processing is restricted to the portion of the right superior temporal cortex that is preferentially recruited by auditory motion in hearing individuals (Battal et al., 2019). Further, we observed that the visually reorganized STC of deaf people is capable of accurately discriminating between different categories of visual motion (i.e. radial, horizontal and stochastic motion), despite not showing selective response to the coherence of visual motion at the univariate level. Importantly, the enhanced decoding for visual motion trajectories in the reorganized portion of the right STC is paralleled by a reduced accuracy of auditory motion decoding in hMT+/V5 in deaf individuals.

Our findings suggest that that the auditory motion-selective portion of the right STC, anatomically overlapping with the *planum temporale*, maintains its functional specialization for motion but redirects its main modality tuning towards visual stimuli following early auditory deprivation, as previously suggested in animal models of congenital deafness (Lomber et al., 2010; Meredith et al., 2011). These findings are also in line with previous studies in early blind individuals reporting functionally specific cross-modal reorganization for auditory spatial (Collignon et al., 2011, 2007) and auditory motion processing (Dormal et al., 2016; Jiang et al., 2016; Poirier et al., 2006) within brain regions in the right hemisphere that, in sighted individuals, preferentially process such stimuli attributes in the visual modality. Our observation also suggests a prominent lateralization of the reorganization process for moving stimuli in the right hemisphere of deaf individuals, consistent with the well-known right lateralization of visuo-spatial processing (Corbetta and Shulman, 2002) and the observation that cortical thickness of the right *planum temporale* is greater in deaf individuals with better visual motion detection thresholds (Shiell et al., 2016). Furthermore, the enhanced multivoxel decoding accuracy, detected in the reorganized right *planum temporale* of deaf people suggests that this region might be capable of representing discrete motion trajectories, a feature that is typically found in occipital region selective for visual motion like hMT+/V5 (Kamitani and Tong, 2006; Rezk et al., 2020).

This adds on exciting prospects in which the development of mind and brain components may not primarily be defined by the sensory modality of the information they operate on, but rather by the computational structure of the problems they solve. The observation that visual motion processing colonizes a functionally related region of the hearing brain also evidences the intrinsic constraints imposed to cross-modal plasticity following early and profound hearing loss. Whether and to what extent developmental trajectories of sensory deprivation modulate the expression of selective cross-modal plasticity remains, however, poorly understood and cannot be evaluated based on the existing studies that investigated early and late deafness separately.

### 4.2. Functional response for visual radial motion in the right A-radial-STC of deaf subjects

In the deaf group, we additionally detected a selective response to visual radial motion in the middle part of the right STC that specifically processes auditory looming motion in the hearing groups; however, no functional preference was observed for horizontal visual motion in any other portion of STC. Since a dynamic staircase procedure was implemented during brain data acquisition, it is unlikely that the radial selectivity found in the STC reflects differences in performance or attention loads across conditions. Behavioural evidence suggests that early deaf individuals do not differ from hearing controls when movement is to be localized in the horizontal plane; instead, they are both faster and more accurate when detecting subtle changes from horizontal to diagonal motion (Hauthal et al., 2013). Moreover, previous investigations have reported a perceptual priority for looming sounds compared to static and horizontal sounds in both human (Moore and King, 1999; Neuhoff, 1998) and non-human primates (Ghazanfar et al., 2002) suggesting predictive strategies to avoid impact or to anticipate motor responses. Similar observations come from studies on the visual system, in which preferential neural patterns for processing of inward radially moving dots, as opposed to outward, have been reported in the human cortex (Jong et al., 1994). Overall, these observations converge on the notion that looming biases might represent an adaptive, increased salience towards ethologically relevant moving objects (Hall and Moore, 2003; Seifritz et al., 2002). There is also evidence that multisensory audio-visual looming stimuli are selectively integrated by humans to facilitate behaviour (Cappe et al., 2009). The neural architecture sustaining multisensory integration of in-depth motion (Cappe et al., 2009; Soto-Faraco et al., 2004) and the environmental salience of approaching/receding stimuli might, therefore, provide the ground for radial-specific visual recruitment of the A-radial-STC in deaf individuals.

It may also be the case that radial, compared to horizontal, visual motion shares more perceptual attributes with speech-related mouth movements (such as protrusions and rounding; Zmarich and Caldognetto, 2003) and speech time-varying acoustic forms that together contributes to shape abstract speech representations in the human brain (Eberhardt et al., 2014). In fact, there is initial evidence that the visual cortex, when early deprived of visual inputs, shows even enhanced functional synchronicity to the acoustic temporal dynamics of speech in blind humans (Van Ackeren et al., 2018), suggesting that early visual and auditory regions may both contributes to the formation of abstracted speech representations. This view is also supported by the finding that speaking faces activates both superior temporal and hMT+/V5 areas in hearing individuals (Paulesu et al., 2003).

### 4.4. Cross-modal reorganization of long-range cortical connectivity and large-scale computational reallocation

What are the mechanisms underlying functional reorganization processes? To gain further insights on this question we conducted a comprehensive investigation on how visual motion information flows between motion-sensitive regions, in the deaf brain, by using dynamic causal modelling. We observed that deaf individuals showed enhanced effective connectivity between the reorganized right A-motion-STC and hMT+/V5, as well as between the right A-motion-STC and IPS, during visual motion perception. Overall these findings are in line with the notion that visual motion information may access the reorganized portions of the *planum temporale* via cortico-cortical connections (Bavelier and Neville, 2002; Gurtubay-Antolin et al., 2020; Rauschecker, 1995). Our work also lends support to previous research showing increased functional coupling between reorganized and preserved sensory cortices sustaining a common specialised function - here motion processing (Benetti et al., 2017; Collignon et al., 2013; Dormal et al., 2016; Shiell et al., 2014). The present findings add to previous reports by showing that, in deaf people, the reorganized right *planum temporale* is part of the brain network dedicated to visual motion perception and that its contribution remarkably resembles that of hMT+/V5 in terms of excitatory/inhibitory interregional dynamics within the motion system. Interestingly, however, the ‘deaf’ A-motion-STC was found to exert a stronger excitatory influence over motion-selective activity in hMT+/V5 compared to its reciprocal from hMT+/V5. A tentative interpretation of this observation might relate to the intrinsic multifaceted representation of motion signals in hMT+/V5 (Rezk et al., 2020) and its potential role as a higher-level hub for motion processing, which might result into higher sensitivity of this region to motion information conveyed by auditory and visual motion-sensitive regions. More specifically, predetermined excitatory afferent connections from A-motion-STC to hMT+/V5 might exceed in quantity and excitatory strength their reciprocal from hMT+/V5 and represent a privileged target for cross-modal recycling of long-range connectivity following early auditory deprivation.

Notably, increased functional connectivity between the hMT+/V5 and the STC during auditory motion has been recently reported in early blind compared to sighted individuals (Dormal et al., 2016), supporting the hypothesis that intrinsic occipito-temporal connections underlying audio-visual integration may play a prominent role in driving the functionally selective cross-modal reorganization observed in case of auditory or visual deprivation (Benetti et al., 2017; Hannagan et al., 2015; Mattioni et al., 2020). Indeed, since motion processing engages both vision and audition, it can be hypothesized that a privileged and intrinsic link already exists between auditory and visual motion selective regions in hearing people (Gurtubay-Antolin et al., 2020; Poirier et al., 2005; Rezk et al., 2020), leading to the emergence of selective cross-modal recruitment in the auditory cortex of early deaf individuals (Dormal and Collignon, 2011; Lomber et al., 2010; Ricciardi et al., 2014). Recently, we also supported this hypothesis in deaf humans by showing that face-selective cross-modal recruitment of the ‘deaf’ temporal voice area, and the large-scale functional reorganization within the face-voice systems, are associated with overall preserved macrostructural connections between visual face- and auditory voice-sensitive brain regions (Benetti et al., 2018).

These findings raise a further compelling question as to how the enhancement of the computational role of temporal area in visual motion processing impacts on the regions typically dedicated to this process in the hearing brain, like hMT+/V5. Interestingly, we observed a significant interaction between group and region in the decoding accuracy of the three motion conditions, revealing that the enhanced decoding observed in the motion-selective temporal region of the deaf is concomitant with a reduction of decoding accuracy in hMT+/V5 when compared to hearing people. This result seems to parallel the observation of higher classification accuracy across auditory motion conditions in the hMT+/V5 of early blind, but higher decoding in the *planum temporale* of the sighted group (Dormal et al., 2016; Jiang *et al*., 2014). In deaf individuals, strengthened occipito-temporal interactions might support a large-scale reallocation of computational resources from spared visual regions to the deafferented auditory cortex, as previously suggested in both deaf and blind individuals (Benetti et al., 2017; Collignon et al., 2013; Klinge et al., 2010; Shiell et al., 2014; but see Bottari *et al*., 2015). It remains a future challenge to understand which computational steps are implemented in the reorganized temporal region of deaf people and their relationship with those implemented in the occipital regions typically implementing visual motion (*e.g*. hMT+/V5).

## Abbreviations

SL: Sign Language
HM: Horizontal Motion
RM: Radial Motion
SM: Stochastic Motion
RDK: Random-Dot Kinematogram
ILD: Interaural Level Difference
STS: Superior Temporal Sulcus
STC: Superior Temporal Cortex
IPS: Intraparietal Sulcus
MVPA: Multivariate Pattern Analysis
LDA: Linear Discriminant Analysis
DCM: Dynamic Causal Model/Modelling
BMS: Bayesian model selection
PEB: Parametric Empirical Bayes

## CRediT author statement

**Stefania Benetti:** Conceptualization, Software, Investigation, Project Administration, Formal Analysis (Behavioural data, fMRI/DCM) Data Curation, Visualization, Writing – Original draft; **Joshua Zonca:** Software, Investigation, Formal Analysis (fMRI); **Ambra Ferrari:** Software, Investigation; **Giuseppe Rabini:** Investigation; **Mohamed Rezk:** Formal Analysis (MVPA); **Olivier Collignon:** Conceptualization, Supervision, Funding acquisition, Writing – Reviewing and Editing.

## Acknowledgments

We thank all the deaf and hearing people who participated in this research for their collaboration. We thank Valentina Foa and Francesca Baruffaldi for their support with subject recruitment and testing, Giulia Dormal for support on the auditory motion data in hearing controls, Emanuele Olivetti (Fondazione Bruno Kessler) for assistance with programming of visual stimuli presentation and Christoph Huber-Huber for guidance on the Bayesian analyses. O.C. is a research associate at the Fond National de la Recherche Scientifique of Belgium (FRS-FNRS).

