## Supplementary Material for "Visual motion processing recruits regions selective for auditory motion in early deaf individuals"

**TITLE**

### Supplementary Material

#### 1. Material and methods

##### *1.1. Auditory Experiment: sound stimuli*

Auditory stimuli consisted of pink noise sounds from 3 different cate-gories: (1) radial (in-depth) motion, (2) lateral motion, and (3) stationary sounds (Fig. 1A). In line with previous neuroimaging studies (Griffiths *et al.*, 2000; Warren *et al.*, 2002; Saenz *et al.*, 2008; Alink *et al.*, 2012; von Saldern and Noppeney, 2013; Dormal *et al.*, 2016), broadband pink noise sounds (44.1 Hz sampling rate) were used since they match the spectrum of most commonly heard frequencies in everyday life without relating to a specific object. Additionally, pink noise minimizes the possibility that a putative occipital response in sighted subjects is a consequence of visual imagery. Moreover, pilot experiments in the scanner revealed that pink noises provided a more vivid sensation of motion relative to pure tones. Sounds lasted either 1 s (standard) or 1.8 s (target)in duration. In the radial motion condition, sounds (mono) either arose or decreased exponentially in intensity (from 10% to maximal intensity and from maximal intensity to 10% intensity) creating the vivid perception of a sound moving toward or away from the listener. In the lateral motion condition, the same sounds were presented separately to the left and to the right ear (stereo) with intensity increasing in one ear while decreasing simultaneously in the other ear, creating the vivid perception of a sound moving from one ear to the other in the azimuth. All participants reported a strong sensation of motion. In the static condition,1 s and 1.8 s pink noise sounds (mono) of constant intensity were presented. A 25 ms ascending/descending ramp was applied at the beginning/end of the static sounds. In order to ensure equal global acoustic energy across conditions despite the application of a ramp in the static condition, the static sounds were normalized based on the mean Root Mean Square (RMS) of the sounds from the motion conditions. Examples of the auditory stimuli used in the present study are provided in the Supplementary Material in Dormal et al., 2016. A block design was implemented in a single run consisting of 30 consecutive blocks (10 repetitions/category) separated by rest periods of9 s. The three categories repeated consecutively with no randomization (i.e. lateral–radial–static). Each block included 18 consecutive auditory stimuli (no ISI) (Fig. 1A). Stimuli within the motion blocks always alternated between the two opposite directions (approaching and receding in the radial condition, left-to-right and right-to-left in the lateral motion condition). The task consisted of detecting longer (1.8 s) sounds by pressing the response button with the index finger of the right hand. Subjects were asked to respond as accurately as possible. Response speed was not emphasized. Within each category, there were 4 blocks with one target (18.8 s duration), 4 blocks with 2 targets (19.6 s duration) and 2 blocks with 3 targets (20.4 s duration). The whole run contained a total of 18 targets *per* category.

*1.2. Auditory motion experiment: fMRI data acquisition*

Functional images were acquired on a 3T TRIO Tim system (Siemens, Erlangen, Germany) equipped with a 12-channel head coil. Multislice T2*-weighted volumes were obtained with a gradient echo-planar sequence applied to the axial plane and acquired in feet-to-head direction with the following sequence parameters: TR = 3.2 mm slice thickness, 0.8 mm inter-slice gap, FoV = 192 x 192 $mm^2$, matrix size = 64 x 64 x 35, voxel size = 3 x 3 x 3.2 $mm^3$ . The 4 initial scans were discarded to allow for steady state magnetization. A structural T1-weighted 3D MP-RAGE sequence was also acquired for all participants with the following parameters: voxel size = 1 x 1 x 1.2 $mm^3$; matrix size = 240 x 256; TR = 2300 ms, TE = 2.91ms, TI = 900ms; FoV = 256; 160 slices.

*1.3. MVPA: Non-parametric statistical analysis*

Multi-class decoding on normally distributed random data collected from small samples (<150 observations) can lead to decoding accuracies that overshoot the chance level merely by chance (Combrisson and Jerbi, 2015). It has been shown that the use of binomial cumulative distributions or non-parametric assessments can better tackle this potential caveat by providing statistical significance levels (*p-values)* that take sample size into account (Combrisson and Jerbi, 2015). Although we included more than 300 trials for each subject entered in the MVPA and in the subsequent parametric assessment, we also implemented an additional non-parametric assessment and applied a binomial formula to further evaluate the significance of the estimated decoding accuracies.

*Motion-decoding within-group*. Statistical significance was assessed using a non-parametric technique by combining permutations and bootstrapping (Stelzer, 2013). For each subject in each ROI, the labels of the different motion conditions were permuted, and the same classification analysis was performed. The previous step was repeated 100 times within each subject. A bootstrap procedure was applied to obtain a group-level null distribution. From each subject's null distribution, one value was randomly chosen (with replacement) and averaged across all participants. This step was repeated 100,000 times resulting in a group-level null distribution of 100,000 values. The statistical significance was estimated by comparing the observed result to the group-level null distribution. This was done by calculating the proportion of observations in the null distribution that had a classification accuracy higher than the one obtained in the real test.

*Motion-decoding between-group.* Statistical significance was estimated non-parametrically using two-sample two-sided t-test. First, in each ROI, a standard two-sample two-sided t-test was performed to compare the two groups. Secondly, a permutation scheme was implemented where the label of the group that subjects belonged to was shuffled. A two-sample two-test was performed on the shuffled data. This permutation scheme was implemented 100,000 times resulting in a null distribution of 100,000 values. The results obtained from the unshuffled data were then compared to the null distribution. The p-values were estimated as the number of absolute t-values from the shuffled data that are higher than the absolute observed t-value from the unshuffled data.

*Statistical significance of classification using a binomial cumulative distribution.* We used a binomial formula, provided by Combrisson and Jaerbi (2015), to estimate the minimal thresholds of decoding accuracy values as a function of selected sample size, class number and expected significance level. This formula, implemented in MATLAB (Mathworks Inc, MA, USA) uses the *binoinv* function to compute the statistically significant threshold $St(\alpha)=\text{binoinv}(1-\alpha, n, 1/c)\times 100/n$, where $n$ is the number of trials, $c$ is the number of classes and $\alpha$ is the significance level given by $\alpha = z/n$ (i.e. the ratio of tolerated false positives $z$ , namely the number of observations correctly classified by chance with respect to all observations $n$).

##### *1.4. Dynamic Causal Modelling: selection of ROIs and time-series extraction*

In the right hemisphere of deaf individuals, each region of interest was first defined as a sphere (5mm radius) centred individually at the local activation maximum closest to the peak of the group-maxima for visual motion selective responses in: (i) the reorganized A-motion-STC (MNI coordinates [60 -34 11]), (ii) the hMT+/V5 (MNI coordinates [45 -64 2]) and the inferior parietal sulcus (IPS, MNI coordinates [48 -34 50]). Then, correspondent time-series were obtained by extracting the first principal component from all raw voxel time series within each region, mean-corrected and high-pass filtered to remove low-frequency signal drifts.

##### *1.5. Dynamic Causal Modelling: model families and models definition*

After defining the two families' features (i.e., target region of the visual motion inputs and modulatory effects of coherent motion on hMT+/V5 and A-motion-STC connectivity), we operationalized the model space based on the factorial combination of the feature levels. Each DCM model would, therefore, be assigned to one of the following families: (1) visual motion input to hMT+/V5 only and no modulation of hMT+/V5-STC connectivity, (2) visual motion input to hMT+/V5 only and modulation of hMT+/V5-STC connectivity, (3) visual motion input to both hMT+/V5 and motion-STC but no modulation of the connectivity between these two regions, and (4) visual motion input to both hMT+/V5 and motion-STC with modulation of the connectivity between these two regions. The stimulus function, which encoded the presence of both coherent and incoherent global visual motion, entered the DCM either through the hMT+/V5 or both this region and the reorganized A-motion-STC. The resulting perturbation was then allowed to propagate throughout the model via 'all-to-all' endogenous extrinsic connections (Friston *et al.*, 2003) between the network nodes. The forward and backward endogenous connections were estimated for each subject independently. The modulatory terms were then modelled based on the family each model would belong to (Figure 6A in main text); while direct modulation of hMT+/V5 connectivity were or were not included depending on the belonging family, modulation of connections from either hMT+/V5 or A-motion-STC, or both, to the higher-level region in IPS were always included. In addition, for each model including forward modulation only we modelled a reciprocal model including both forward/backward modulation. After Bayesian model inversion, approximation of the log model evidence for each model was used to implement family and model selection. We used a two-folded partitioning approach in which families (1)/(2) were compared to families(3)/(4) and models within each group of families averaged. This process yielded the selection of families (3) and (4), which were subsequently compared against each other. Subsequently, the optimal model for the deaf group was identified within family 3 and 4 by applying random-effects Bayesian Model Selection (as implemented in SPM12; Stephan *et al.*, 2009), which selects the model that offers the most accurate and least complex explanation for the data (i.e., the highest free energy/log model evidence).

*1.6. Dynamic Causal Modelling: Parametric Empirical Bayes – automated search*

In DCM, a model represents a set of differential equations that describe how experimental stimulation translates into observed data through neural activity. Each model has connectivity parameters, which are estimated from the observed data using a Bayesian probabilistic inversion method called Variational Laplace (Friston *et al.*, 2007) and can be used to quantify the commonalities (or differences) across individuals or groups. To this purpose, DCM is supplemented with a hierarchical model with random effects on parameters (rather than models), the Parametric Empirical Bayes (PEB) framework (Friston *et al.*, 2016). In this framework, after model specification and inversion, the individual parameters of interest are collated and modelled at the group-level using a General Linear Model (GLM) that includes both the expected values and the uncertainty (covariance) of the estimated parameters (for the description of PEB useful implications see Zeidman et al., 2019).

In this study, we implemented PEB with the purpose to characterize, across deaf individuals, a common profile of changes in inter-regional coupling, within the reorganized motion-network, that significantly contributed to visual motion processing (irrespective of motion coherence) in the deaf group. Therefore, we first defined and inverted a 'full' DCM model including visual motion inputs to both hMT+/V5 and A-motion-STC, as well as 'all-to-all' endogenous extrinsic (i.e. independent of experimental manipulation of motion coherence) connection between network regions. Subsequently, we collated the estimated parameters for all subjects and configured a PEB model by specifying the group-level GLM design matrix with the mean experiment-related connectivity changes (constant of the group mean) as a between-subject factor of interest and all the endogenous connections as within-subject factor, enabling visual motion to potentially influence all connections. This PEB model was then inverted resulting in a set of estimated group-level parameters and the group-level free energy, which is the sum of individual DCMs accuracies minus the complexity induced by fitting the DCMs and the group-level GLM. Since we had no strong hypothesis on which endogenous connections would be mostly influenced by visual motion, we adopted an exploratory approach by using Bayesian Model Reduction (BMR) to automatically search over reduced models. For this study, the GLM contained 9 parameters in total and the aim of the automated search was to find the best reduced GLM with certain parameters switched off. The BMR procedure compares the evidence for reduced models and iteratively discharges parameters that do not contribute to model evidence until discarding any parameter starts decreasing model evidence. A Bayesian Model Average (BMA) is then calculated over the models from the final iteration, posterior parameter estimates thresholded based on the free energy criterion of posterior probability >95% and the results reported in a connectivity matrix (Figure 6D-top).

#### 2. Results

Given the relative small size of the deaf sample, we conducted a series of power analyses (as implemented in G*Power software; Faul *et al.*, 2007) on effect size and variance estimates of (1) results published (actual data) in Benetti et al., 2017, and (2) the results reported in Bola et al., 2017 (estimates obtained from the graphical bar plots reported and data information reported in the published article). Since these datasets refer to experimental conditions that differ from those tested in the present work we decided to adopt an even more conservative approach and estimate the expected power given a required alpha-error probability of p=0.01, instead of p =0.05. Additionally, since we are well aware of the implicit limitations of post-hoc power analyses implemented on published data (see Hayasaka *et al.*, 2007; Mumford, 2012), we also complemented these power analyses with (1) a sensitivity analysis (as implemented in G*Power software) and (2) a Bayes factor estimation to test the relative predictive performance of competing hypotheses and plausibility of the parameter values (JASP Version 0.13.1, JASP Team (2020); Wagenmakers *et al.*, 2018b, a) on the present dataset.

*2.1. Post-hoc Power analyses on previously published data*

*2.1.1 Interaction between stimuli (faces,houses) and group (hearing, hearing signers, deaf) in Benetti et al., 2017.* The estimated power to detect a significant interaction between visual stimuli and experimental group (i.e. face selectivity in the deaf sample) is 0.960 (critical F=2.404, d.f.=10) given an estimated effect size of F(V)=0.866 (partial $n^2 = 0.429$), an alpha error of p=0.01 and a total number of 46 (16 hearing, 15 hearing SL signers, 15 deaf) subjects.

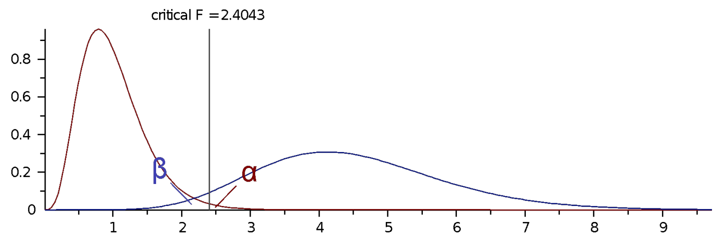


Based on the above calculations, we estimated that the hypothetical sample size required to observe similar findings of selectivity is 33 subjects in total across the three groups (i.e. 11 subjects per group) at a power value of 0.80 and alpha-error probability of p=0.01 and given an effect size of 0.866. We also estimated the required total sample size for a range of given effect sizes spanning from 0.6 to 1.2 (reported in the plot below): if we assume a total sample size of 39 (as in the present study) subjects (13 in each experimental group) with a power value of 80% we can detect true differences as small as f(V)= 0.839 and above.

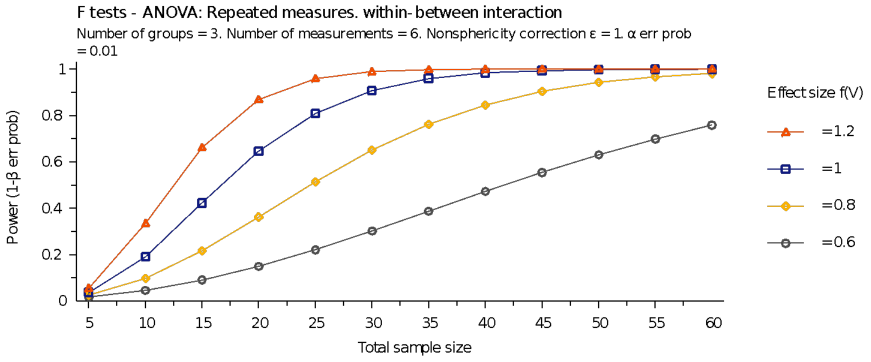


*2.1.2 Main effect of group (deaf vs. hearing) for faces-houses as in Benetti et al., 2017.* The estimated power to detect significant differences between deaf and hearing (non-signers) is 0.930 (critical t=2.462, d.f. = 29) given an effect size of Cohen's d= 1.436, an alpha error of 0.01 and sample sizes of 16 (hearing) and 15 (deaf).

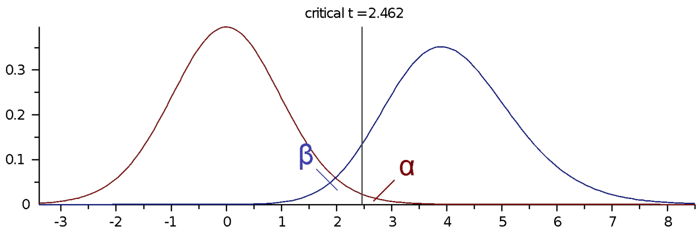


Based on the above calculations, we estimated that the hypothetical total sample size required to observe similar findings of selectivity is 24 subjects in total across the three groups (i.e. 12 subjects per group) at a power value of 0.80 and alpha-error probability of p=0.01 and given an effect size of 1.436. We also estimated the required total sample size for a range of given effect sizes spanning from 0.6 to 1.4 (reported in the plot below): <u>if we assume a total sample size of 26 subjects (13 in each experimental group) a power value of 80% is hypothesised for an effect sizes in the range of(V)= 1.2-1.4 and above.</u>

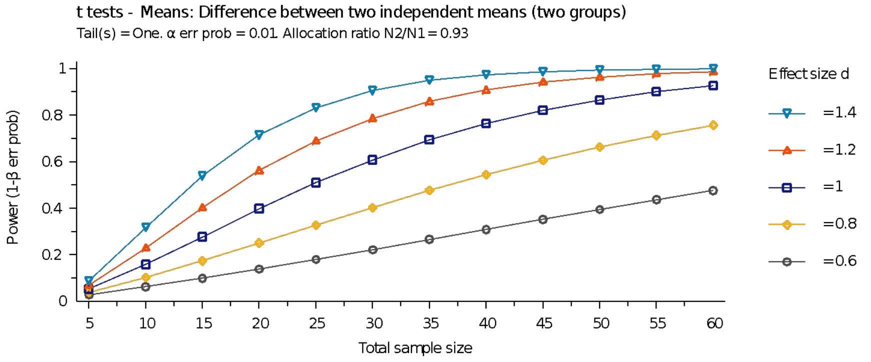

*2.1.3 Main effect of group (deaf vs. hearing signers) for faces-houses as in Benetti et al., 2017.* The estimated power to detect face selectivity in the deaf compared to the hearing signers is 0.917 (critical t=2.467, d.f.=28) given an effect size of Cohen's d=1.427, an alpha error of 0.01 and a sample size of 15 individuals in each experimental group.

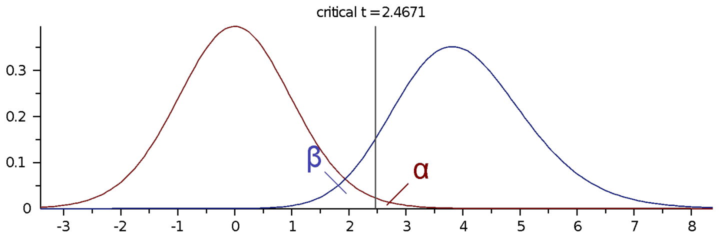


Based on the above calculations, we estimated that the hypothetical total sample size required to observe similar findings of selectivity is 24 subjects in total across the three groups (i.e. 12 subjects per group) at a power value of 0.80 and alpha-error probability of p=0.01 and given an effect size of 1.4.

*2.1.4 Main effect of group (deaf vs. hearing) for visual rhythm-visual control as in Bola et al., 2017.*

No estimates of the actual effect size or of the variance within the groups was reported in the published article. We therefore calculate the hypothetical effect size and power based on a conservative approximation of estimated group mean betas for [visual rhythm-visual control] (deaf = 0.60 and hearing = 0.05) and the reported SEM (deaf = 0.12 SEM/0.464s.d.; hearing = 0.11 SEM/0.425 s.d.) and sample sizes (15 in each group) as reported in the bar plot and Methods section. The estimated power to detect significant differences between deaf and hearing is 0.814 (critical t=2.467, d.f. = 28) given an estimated effect size of Cohen's d= 1.236, and an alpha error of 0.01.

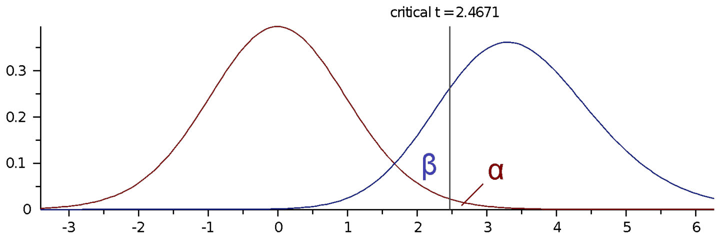


Based on the above calculations, we estimated that the hypothetical total sample size required to observe similar findings of selectivity is 30 subjects in total across the two groups (i.e. 15 subjects per group) at a power value of 0.80 and alpha-error probability of p=0.01 and given an effect size of

1.2. If we assume a total sample size of 26 subjects (13/13 in each experimental group) a power value of 80% is hypothesised for an effect sizes as large as d= 1.267 and above. If the alpha error is set to p=0.05 the detectable effect size drops to a value of d=0.967.

*2.2 Sensitivity analyses on the main results of the present study*

*2.2.1 Main effect of group (Deaf vs. Hearing-nSL conj.Hearing-SL) for global visual motion in the A-Motion-STC area* . We implemented a sensitivity analysis on the present sample of participants. This reveal that with a total sample of 42 subjects (15 Hearing-nSL, 14 Hearing-SL and 13 Deaf) a required power of 80% and an alpha-error of p=0.01 a difference between groups can be detected that is as small as F(V)=0.610 in a one-way ANOVA. With an alpha error of p=0.05, the minimum detectable effect size would be F(V)=0.497. The estimated effect size for the observed main effect of group is $\eta_p^2 = 0.310 = 0.670$ f(V). The estimated effect sizes for the 2 post-hoc comparisons are as follows: t(deaf > hearing-nSL) = 3.425, $P < 0.002$; Cohen's d = 1.298; t(deaf > hearing-SL) = 3.771, $P < 0.001$, Cohen's d = 1.452.

2.2.2 *Interaction between stimuli (horizontal vs. radial visual motion) and groups in the A-radial-STC area.* This analysis revealed that with a total sample of 42 subjects (15 Hearing-nSL, 14 Hearing-SL and 13 Deaf), 9 different measurements (3 motion condition by 3 groups), a required power of 80% and an alpha-error of p=0.01 a significant interaction can be detected that is as small as F(V)=0.807 in a repeated measures ANOVA. With an alpha error of p=0.05, the minimum detectable effect size would be F(V)=0.691. The estimated effect size for the observed interaction (= radial motion > horizontal motion in Deaf compared to Hearing individuals) is $\eta_p^2 = 0.440 = 0.899$ f(V). The estimated effect size for the two post-hoc ANOVAs are as follows: F(radial, deaf > hearing) = 9.841, $p < 0.001$, $\eta_p^2 = 0.327 = 0.697$ F(V) ; F(horizontal, deaf > hearing) = 1.873, P = 0.167.

*2.3.1 Bayesian ANOVA on global visual motion in the A-Motion-STC area.* This analysis revealed that the data collected in this ROI were 44.285 times more likely to occur under the model including the effect of the group, compared to the model without the effect. Post-hoc comparisons of Deaf *vs.* Hearing-nSL and Deaf *vs.* Hearing-SL revealed posterior odds of 10.16 and 19.79 respectively, which indicates strong evidence in favour of the alternative hypothesis of visual cross-modal recruitment of this region during visual global motion processing in deaf compared to hearing individuals.

*2.3.2 Mixed-factor Bayesian ANOVA on radial and horizontal visual motion in the A-radial-STC area.* This analysis determined that the data collected in this ROI were best explained by a model that included both main factors, the Group and Motion direction, and their interaction. The Bayes factor ($BF_{10}$) for the model was 1901.88, while $BF_{10}$ = 1174.68 for the interaction term and on top of the main effects. These estimates indicate strong evidence in favour of this model compared to the null model. A series of post-hoc comparisons reported adjusted posterior that provide strong evidence for a difference between Deaf *vs.* Hearing-nSL (= 215.67) and between Deaf *vs.* Hearing-SL (= 719.07) for mean weight values of [Radial – Horizontal] in this region.

*2.4 ROI multivariate pattern analysis: non-parametric assessment*

*2.4.1 Permutation and bootstrapping*

We tested whether the classification of the three motion categories (i.e. HM, RM, SM) in the right A-motion-STC and hMT+/V5 and in the two experimental groups (i.e. ED and H-$nSL_{conj}$.H-SL) was above chance-level (33.3%). Non parametric significance testing (false discovery rate (FDR)-corrected) confirmed that visual motion-category decoding was above chance in the right hMT+/V5 of both hearing ($p = 0.0004$) and deaf ($p = 0.0008$) individuals but only in the right A-motion-STC of deaf participants ($p = 0.0011$ in ED; $p = 0.1697$ in H-$nSL_{conj}$.H-SL).

*2.4.2 Significance thresholds of decoding accuracy: binomial formula*

The binomial formula applied to our dataset revealed that, given 360 trials for each subject included in the MVPA and an expected false-positive ratio $p < 0.0125$ (corrected for 4 multiple comparisons), a 3-class decoding accuracy is statistically significant only if it exceeds the threshold of 38.8%. This additional assessment confirmed the significance of the decoding accuracy values in the right hMT+/V5 of both experimental groups (ED = 41.89%, 95% CI[36.27,47.52]; Hearing = 42.65%, 95% CI [42.60,48.00]) but only for the deaf individuals in the right A-motion-STC (ED = 40.97%, 95% CI[36.17,45.76]; Hearing = 34.86%, 95% CI[31.85,37.88]).

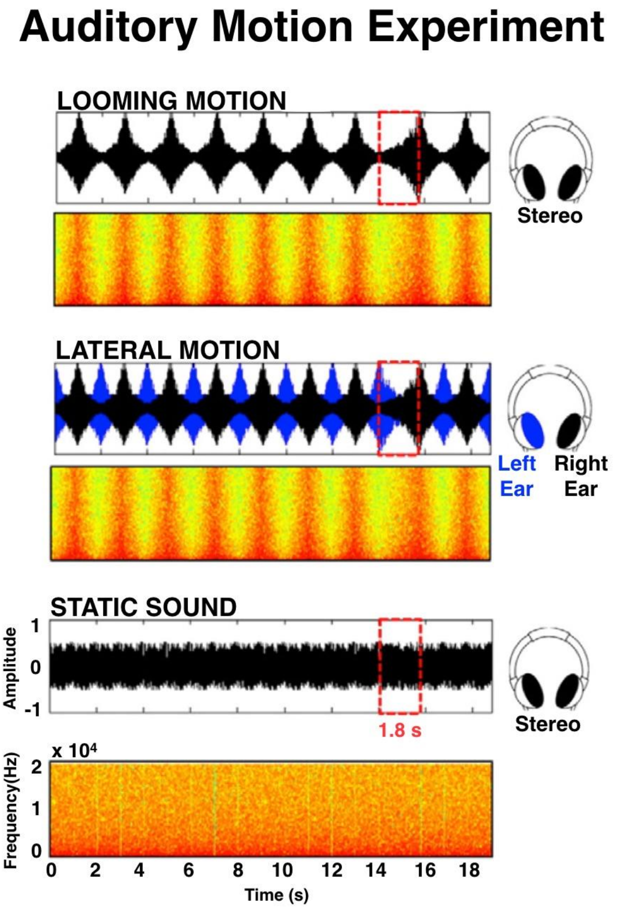


**Figure S1.** Graphical depiction of the stimuli and the procedures used in the auditory motion experiment in hearing-nSL individuals. Acoustic properties are illustrated for representative blocks of each experimental condition. Graphic panels represent the amplitude of a block as a function of time (waveform) and the spectrum of frequencies as a function of time (frequency spectrum). Acoustic properties of a 1.8s target sound are highlighted within red dashed-line boxes. The sounds were delivered in-phase to each ear in the radial and static condition (black waveforms represent sounds delivered to both ears), and out-of-phase in the lateral condition (black/blue waveforms represent the sounds delivered to the right/left ear respectively).

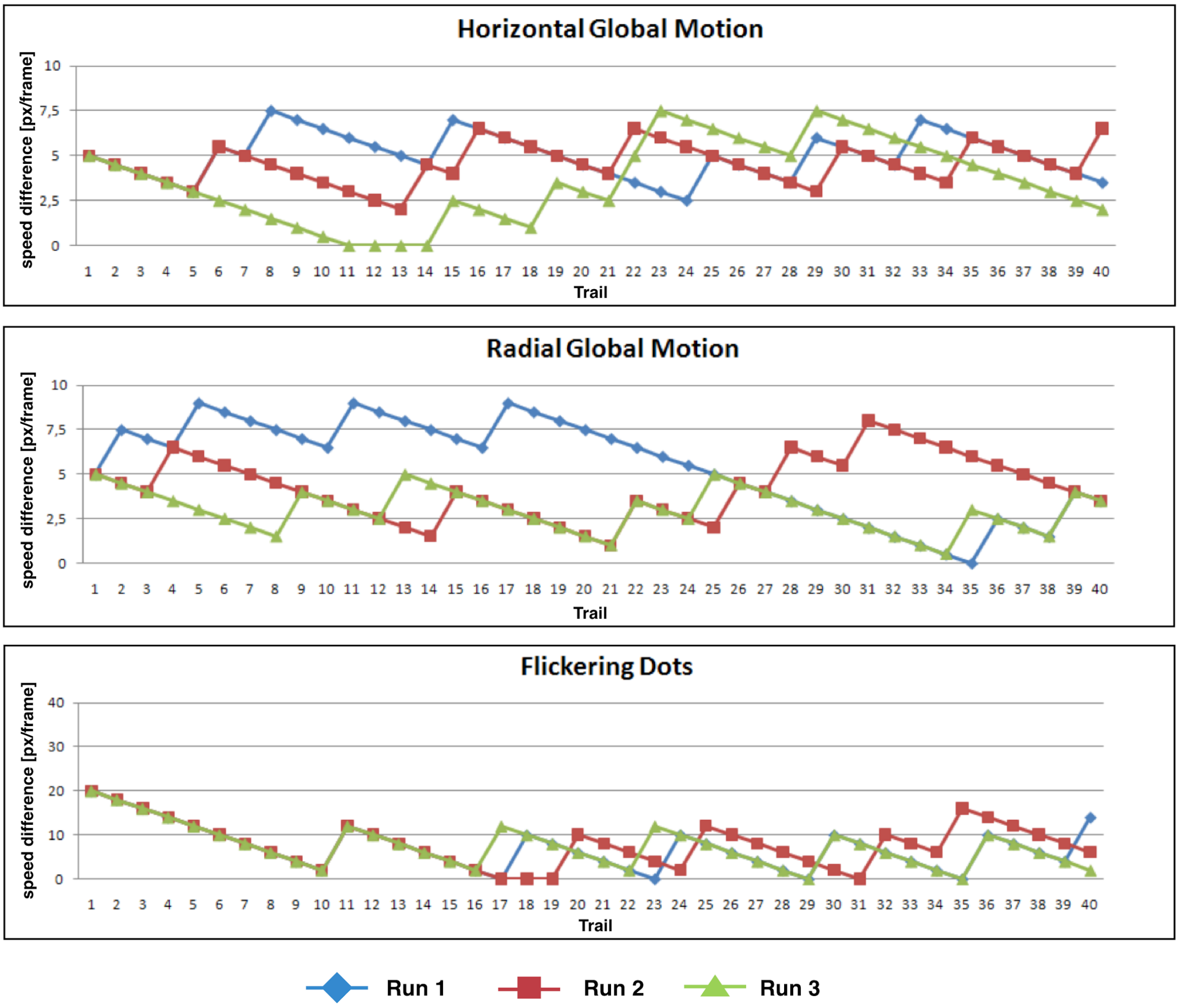


**Figure S2.** Staircase performance profile, in terms of speed difference between moving dot arrays at each trial, of a representative individual. The three experiment runs are coded in different colours. The staircase procedure adjusted the speed gap between each couple of dot arrays 1-step-down for correct response and 5-steps-up for wrong response.

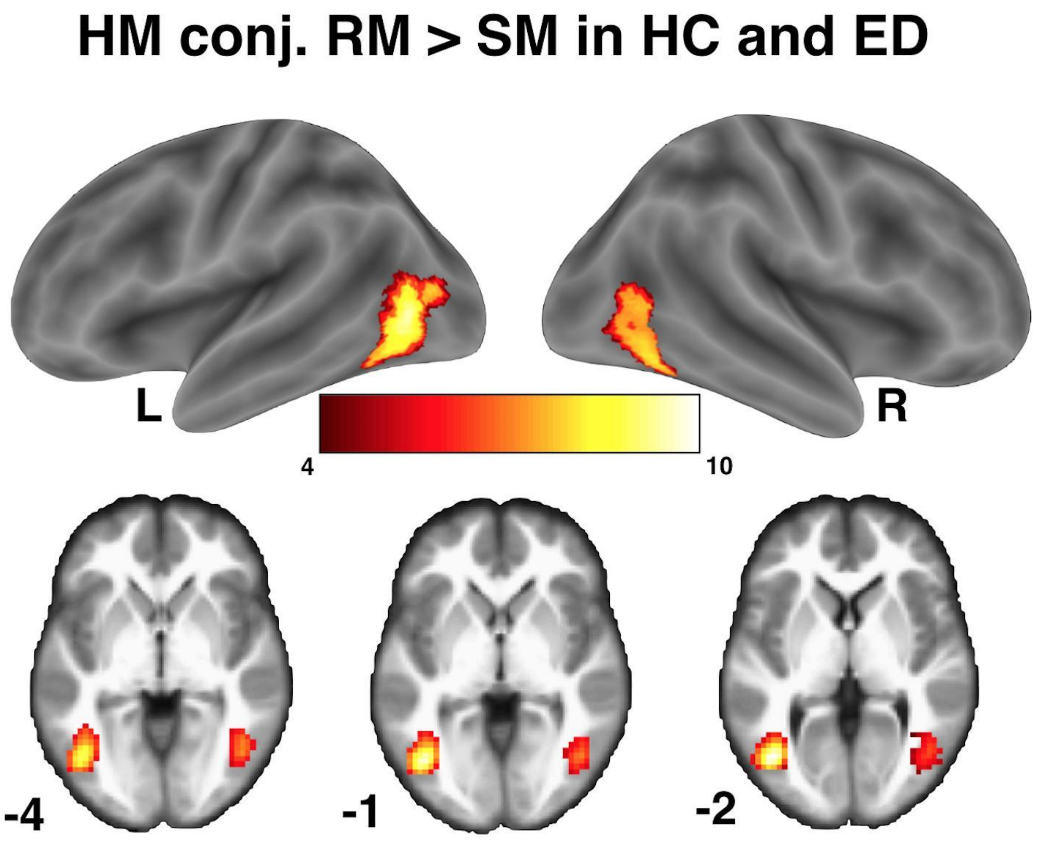


**Figure S3.** Regional responses to coherent *vs.* non-coherent visual motion in the three groups. Supra-threshold effects are depicted at P < 0.05 FWE cluster-corrected with a minimum size of 20 contiguous voxels (160mm$^3$) per cluster and superimposed on MNI renders (BSPMVIEW; http://www.bobspunt.com/bspmview/). The activations shown here for HC (Hearing-nSL and Hearing-SL individuals) and ED (Early deaf individuals) refer to the conjunction analysis across the three experimental groups. Peak-coordinates (MNI) for group-maxima: left [-42 -70 2] and right [45 -64 8]. HM, Horizontal Global Motion; RM, Radial Global Motion; SM, Stochastic Motion; L, Left; R, Right.

**Video S1. Animated representation of the Visual Motion fMRI experiment.** The video illustrates the stimuli used in each of the 3 conditions tested as well as the structure of stimulus delivery. Explanatory subtitles are provided.

**Table S1** Characteristics of the early deaf participants.

| Code | Deafness Onset | Deafness Severity | Deafness Duration | Preferred Language | Hearing Aid Use | Experiment |
|---|---|---|---|---|---|---|
| ED1 | Birth | Profound | 21 | LIS | no | fMRI-MEG |
| ED2 | Birth | Profound | 45 | LIS | Partial | fMRI-MEG |
| ED3 | Age 0-4* | Profound | 32 | LIS/Italian | Full | fMRI-MEG |
| ED4 | Birth | Severe/ Profound | 39 | Italian | Full | fMRI-MEG |
| ED5 | Birth | Profound | 31 | Italian/LIS | Full | fMRI-MEG |
| ED6 | Birth | Profound | 34 | LIS | No | fMRI-MEG |
| ED7 | Birth | Profound | 41 | LIS/Italian | Partial | fMRI-MEG |
| ED8 | Birth | Profound | 31 | LIS | No | fMRI |
| ED9 | Birth | Severe | 24 | Italian/LIS | Full | fMRI |
| ED10 | Birth | Severe | 25 | Italian | Full | fMRI-MEG |
| ED11 | Birth | Profound | 24 | LIS/Italian | Full | fMRI-MEG |
| ED12 | Birth | Profound | 39 | LIS | Full | fMRI-MEG |
| ED13 | Birth | Profound | 36 | LIS/Italian | No | fMRI-MEG |

*Hearing Aid use: Partial = only during school or work hours; Full = on most of the day to support environmental sound detection (alarms, door bells, foot steps). Only ED10 reported support during speech reading. Abbreviations: LIS, Italian Sign Language. *ED3 reported measles before age 4.*

**Table S2.** Group-specific peak-coordinates used for definition of regions of interest in the present study.

| Area | $X_{(mm)}$ | $Y_{(mm)}$ | $Z_{(mm)}$ |
|---|---|---|---|
| *fMRI Auditory Motion Localizer: ROIs definition (9 mm spheres)* | | | |
| Motion-STC | | | |
| Right | 64 | -36 | 14 |
| Left | -56 | -28 | 8 |
| Lateral-STC | | | |
| Right | 48 | -28 | 28 |
| Left | -52 | -26 | 22 |
| Looming-STC | | | |
| Right | 56 | -17 | -2 |
| Left | -56 | -24 | 4 |
| *Multivariate pattern analysis (9mm spheres):* | | | |
| Right motion-STC | 60 | -34 | 11 |
| Right hMT+/V5 | 48 | -67 | 5 |
| *Dynamic Causal Modelling (5mm spheres)* | | | |
| Right motion-STC | 60 | -34 | 11 |
| Right hMT+/V5 | 48 | -67 | 5 |
| Right IPS | 48 | -34 | 50 |

STC = Superior Temporal Cortex; IPS = Intraparietal Sulcus
